# The androgen receptor polyglutamine tract sets the transcriptional balance between muscle growth and atrophy

**DOI:** 10.64898/2026.09.29.755413

**Authors:** Francis N. Grafton, Narjes Rohani, Wooi Fang Lim, Yang Kai Ng, Alessandro Fichera, Salma Ali, Virginia Scappini, Alfina A. Speciale, Rebecca Chalcraft, Antonio Garcia Guerra, Yasumasa Hashimoto, Jennifer Frommer, Maria Pennuto, Esther Garcia Gonzalez, Laura Zanetti-Domingues, Christopher J. Tynan, Stephan J. Sanders, Michele Pansini, Matthew J. A. Wood, Carlo Rinaldi

## Abstract

The androgen receptor (AR) mediates the potent anabolic effects of testosterone and other androgens on skeletal muscle, yet the transcriptional mechanisms coupling this hormonal signal to muscle growth remain unresolved. AR is a ligand-inducible transcription factor whose N-terminal transactivation domain (NTD) harbours a polymorphic polyglutamine (polyQ) tract that, when pathologically expanded, causes spinal and bulbar muscular atrophy (SBMA). To define how the NTD and its polyQ tract shape muscle transcription, we combined an engineered AR allelic series in human myogenic cells with super-resolution and single-molecule imaging, genome-wide binding and expression maps, and human population genetics. Under physiological conditions, androgen-activated AR assembles into dynamic transcriptional condensates and drives an anabolic programme in skeletal muscle through activation of the AP-1 pathway. Progressive expansion of the polyQ tract destabilises AR chromatin binding and shifts its occupancy from promoters to low-affinity distal enhancers, tracking with imaging signatures of muscle aging. In SBMA, this culminates in aberrant transactivation of ectodysplasin A2 receptor (EDA2R), a marker of senescence and inflammation, which engages non-canonical NF-κB signalling to drive muscle atrophy. Silencing Eda2r attenuated muscle wasting in SBMA mice, establishing this axis as therapeutically actionable. Together, these findings reveal a fundamental mechanism of skeletal-muscle homeostasis and open therapeutic avenues for the many conditions in which muscle atrophy is central.

## Introduction

Androgens, the steroid hormones responsible for male sexual development, are among the most potent naturally occurring anabolic agents in humans, increasing muscle fibre size and strength in a dose-dependent manner^1,2^. Low levels of circulating androgens and weak grip strength are strong predictors of mortality across populations^3–5^.

While being widely administered clinically to treat hypogonadism and to counter the muscle loss associated with aging, cancer cachexia, and chronic disease, androgens are also the most frequently misused performance-enhancing drugs worldwide^6,7^, despite being constrained by adverse effects such as erythrocytosis, infertility, and cardiovascular accidents^8,9^. Remarkably, the molecular events underpinning the anabolic effects of testosterone on skeletal muscle remain poorly defined^10^.

The effects of testosterone are mediated by the androgen receptor (AR), a ligand-activated transcription factor (TF) of the nuclear receptor superfamily, whose activity in myofibres is required for androgen-driven gains in muscle mass and strength^11^. Upon androgen binding, AR undergoes conformational changes, translocates to the nucleus and binds chromatin to drive gene expression programmes. How AR transcriptional activity regulates skeletal muscle homeostasis is a central unanswered question. Closing this gap is essential to unravel a fundamental mechanism of physiology and guide the development of safe and effective anabolic therapies.

A powerful conceptual framework for productive transcription has emerged from the recognition that TFs and their binding partners assemble into biomolecular condensates that concentrate the transcriptional machinery at active genes^12^. This capacity is encoded largely by intrinsically disordered regions (IDRs) within transcription factor activation domains, whose multivalent, low-complexity interactions drive the phase separation that underlies gene activation^13^. AR is an archetype of this class: it possesses a large, intrinsically disordered N-terminal domain (NTD) which mediates condensate formation and partitioning with transcriptional effectors, such as the co-activator Mediator (MED1) or RNA polymerase II (RNAPII)^14–17^.

Strikingly, TFs frequently harbour microsatellite repeat expansions in their IDRs, providing a natural genetic handle to interrogate their function^18,19^. AR is no exception: its NTD harbours a microsatellite CAG repeat encoding a polyglutamine (polyQ) tract, which, when expanded beyond the threshold of 38 repeats, causes spinal and bulbar muscular atrophy (SBMA, Kennedy’s disease, OMIM 313200), an X-linked neuromuscular disease characterized by lower motor neuron degeneration and primary skeletal muscle involvement^20,21^.

Together, these features (i.e.: ligand-activatable chromatin binding and disease-causing microsatellite repeat in its intrinsically disordered NTD) make AR a uniquely powerful system for dissecting transcription-factor behaviour. Defining how AR condensate formation, chromatin dynamics and transcriptional output are governed in skeletal muscle therefore promises to reveal not only the molecular basis of androgen anabolism, but also the mechanistic origins of SBMA pathology and, more broadly, how repeat expansions affect transcription factor function.

### The polyglutamine tract regulates AR transcriptional activity

In order to study AR transcriptional activity in a close-to-endogenous setting, we first proceeded to knock-out *AR* by CRISPR-Cas9 in human skeletal muscle cells (hMyo) (**Supplementary Fig. 1a and 1b**). These cells were then engineered to stably express the human AR transgene with increasing polyQ tracts, ranging from wild type (24Q) to pathological (66Q, and 100Q), or a protein variant lacking the amino (N) terminal domain altogether (ΔN), under a tetracycline response element to enable matching levels with endogenous expression (**Supplementary Fig. 1c and 1d**).

**Fig. 1.**
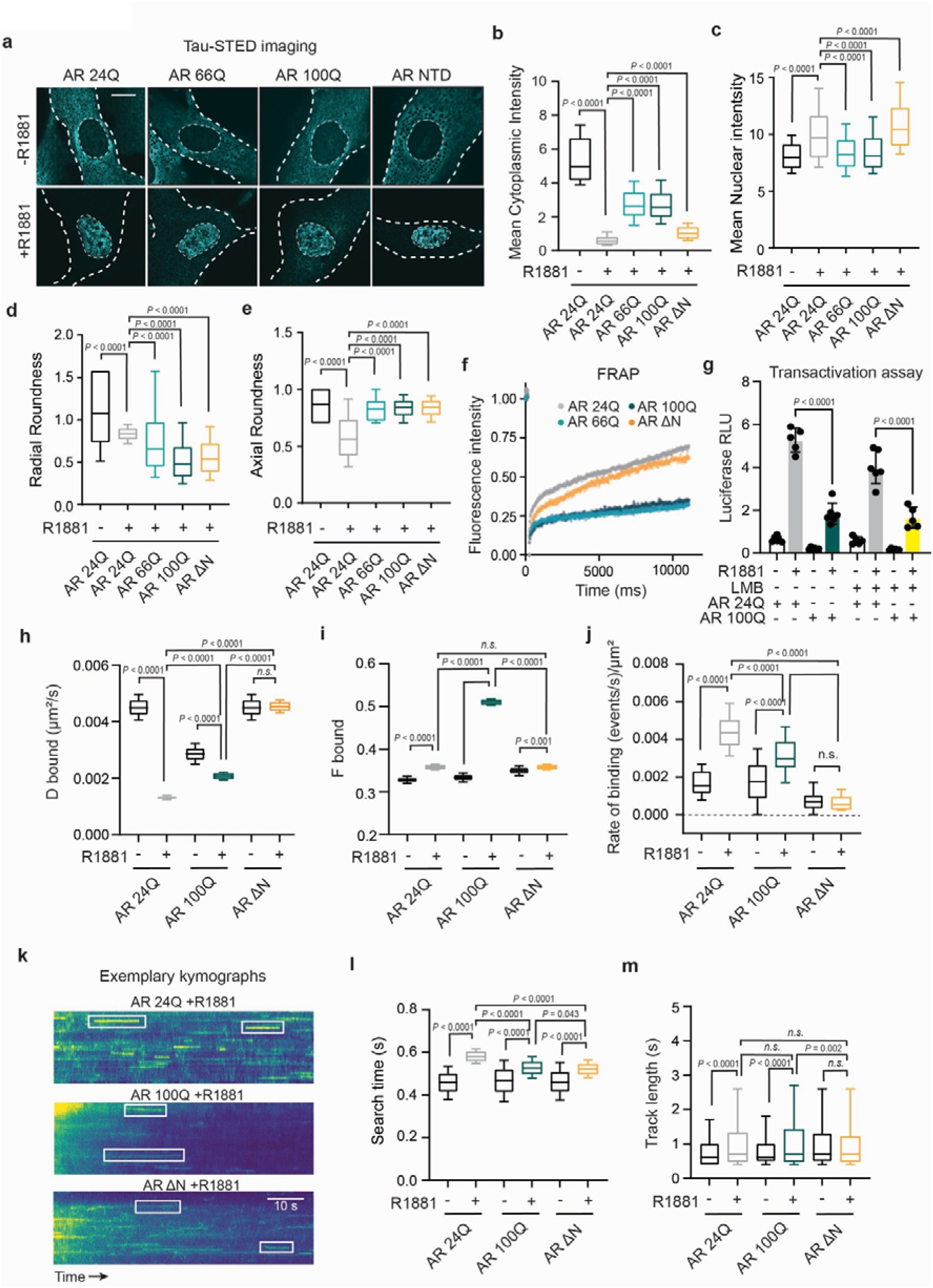
AR polyglutamine tract regulates its transcriptional activity. **a**, Representative stimulated-emission-depletion (Tau-STED) images of HT-tagged AR in human myoblasts (hMyo) expressing AR 24Q, 66Q, 100Q or NTD-deleted AR (AR ΔN), in the absence (top) or presence (bottom) of 10 nM R1881 Dashed lines outline the nuclear and cell boundaries. Scale bar, 20 µm. **b**, **c**, Quantification of mean cytoplasmic (**b**) and mean nuclear (**c**) AR signal intensity across genotypes ± R1881. PolyQ-expanded AR (66Q, 100Q) shows elevated cytoplasmic and reduced nuclear signal relative to AR 24Q upon ligand activation (*n* = 5 cells per replicate, 3 replicates per condition). **d**, **e**, Radial (**d**) and axial (**e**) roundness of AR nuclear condensates ± R1881. AR 100Q and AR ΔN condensates show significantly reduced sphericity relative to ligand-activated AR 24Q (*n* = 5 cells per replicate, 3 replicates per condition). **f**, Fluorescence recovery after photobleaching (FRAP) of AR nuclear condensates for AR 24Q, 66Q, 100Q and AR ΔN (*n* = 3 cells per replicate, 3 replicates per condition). AR 66Q and AR 100Q recover more slowly than AR 24Q and AR ΔN. Data are mean ± s.e.m. One-way ANOVA with Tukey’s multiple-comparisons correction. **g**, AR transactivation measured by luciferase reporter (relative light units, RLU) ± R1881 and ± the nuclear-export inhibitor leptomycin B (LMB) (*n* = 5-6). Enforced nuclear retention does not rescue the transactivation deficit of polyQ-expanded AR. Data are mean ± s.e.m. One-way ANOVA with Tukey’s multiple-comparisons correction. **h**, **i**, **j**, Single-particle tracking (SPT) parameters from a two-state kinetic model ± R1881: diffusion coefficient of the bound state (D bound, **h**), bound fraction (F bound, **i**) and rate of binding events per µm² (**j**). R1881 lowers D bound and raises F bound and the binding rate in AR 24Q; AR 100Q shows a smaller D-bound reduction, a markedly higher F bound and a blunted binding-rate increase; AR ΔN shows a high, ligand-insensitive D bound with minimal F-bound and binding-rate changes (*n* > 10 cells per replicate, 3 replicates per condition). **k**, Exemplary kymographs of single AR molecules for AR 24Q, AR 100Q and AR ΔN, treated with R1881. Each kymograph displays fluorescence intensity along a nuclear region (vertical) as a function of time (horizontal; time increases to the right). Stable binding events appear as sustained horizontal tracks (examples in white boxes). Scale bar, 10 s. **l**, **m**, Search time (**l**) and track length (**m**) from SPT ± R1881. Androgen activation prolongs the search time of AR 24Q but shortens it in AR 100Q and AR ΔN; track length increases with R1881 in AR 24Q and AR 100Q but not in AR ΔN (*n* > 10 cells per replicate, 3 replicates per condition). **b-e**, **h**, **i**, **l**, **m**, Box plots show the median (centre line), interquartile range (box) and 10th–90th percentile (whiskers); one-way ANOVA with Tukey’s multiple-comparisons correction. *P* values are indicated on each panel; *n* represents the number of experiments; n.s., not significant.

Previous studies have highlighted that ligand-activated AR can form transcriptionally active condensates through multivalent interactions mediated by its intrinsically disordered NTD^14,16,17,22–24^. To address the role of the polyQ tract in AR nuclear condensation, we performed Tau-STED microscopy, a powerful super-resolution imaging technique used to resolve the nanoscale architecture of transcriptional condensates by achieving lateral resolutions down to 20-50 nanometers^25^. To this end, we used a set of the hMyo allelic series whereby AR was tagged at the carboxy (C) terminal with HaloTag (HT)^26^, an ideal tool for super resolution microscopy techniques, due to its combined properties of specific covalent binding, uniform brightness, and improved photostability^27,28^ (**Supplementary Fig. 1a and 1d**).

As expected, treatment with synthetic androgen metribolone (R1881) drove AR nuclear translocation (**Fig. 1a and 1b**). In the polyQ-expanded lines (AR 66Q and AR 100Q), a larger fraction of AR molecules was retained in the cytoplasm, resulting in elevated cytoplasmic intensity and reduced nuclear signal (**Fig. 1a and 1b; Supplementary Fig. 2a and b**). Because this deficit was already apparent after 1 h post ligand induction (not shown), we hypothesise that this observation reflects a delay in nuclear import. These findings, together with previously reported impairment of nuclear export of polyQ-expanded AR^29^, support a model of globally impaired AR nucleocytoplasmic dynamics upon polyQ expansion.

Since liquid-like condensates naturally tend towards spherical shapes to minimize surface tension, we next sought to determine how the NTD determines AR condensate morphology. Both polyQ-expanded AR condensates showed significantly reduced sphericity relative to AR 24Q upon ligand activation (**Fig. 1d and 1e**), consistent with a liquid-to-gel transition of the kind observed upon cellular stress or aging^30^. The same loss of sphericity was observed for ΔN AR, indicating that an intact, healthy disordered NTD is required to scaffold well-formed condensates. Reduced sphericity can arise from two opposite states: an overtly rigid condensate, as in polyQ-expanded AR, or an insufficiently scaffolded assembly, as in ΔN AR. To test this interpretation, we performed fluorescence recovery after photobleaching (FRAP)^31^, which confirmed opposite dynamic signatures underlying the shared morphological phenotype: slow recovery for polyQ-expanded AR and rapid for ΔN AR (**Fig. 1f**).

Previous reports from us and others have shown that polyQ AR has reduced transactivation activity compared to wild type^32,33^. In order to exclude the possibility that our observation of increased cytoplasmic retention simply reflects fewer AR molecules reaching the nucleus, we performed a luciferase-based AR transactivation assay in the presence of leptomycin B, a selective inhibitor of nuclear export^34^. Increasing AR nuclear retention was not sufficient to normalise the transactivation deficit of polyQ-expanded AR (**Fig. 1g**), pointing to an intrinsic defect in chromatin engagement rather than a simple deficit in nuclear abundance.

Because transcription is a dynamic, rapid and stochastic process that conventional imaging cannot resolve, we turned to single-particle tracking (SPT) microscopy, to quantify transcription factor dynamics in living cells and at single-molecule resolution^35^. From a two-state kinetic fit of the displacement distributions (**Supplementary Fig. 3a**), we extracted three parameters that report distinct features of transcription-factor binding: (i) the diffusion coefficient of the bound state (D bound), which reflects how tightly captured molecules are held, ii) the fraction of molecules in the bound state (F bound), which reflects how many molecules are captured, and iii) the rate of binding events. Combining these parameters resolved three distinct patterns (**Fig. 1h-j**): Firstly, AR 24Q responded to androgen treatment with reduced D bound and increased F bound and rate of chromatin binding events, indicating that, upon androgens, AR molecules are firmly anchored onto the DNA. Secondly, polyQ-expanded AR showed partial reduction in D bound, a markedly increased F bound, and a blunted increase in rate of binding events, suggesting that activated polyQ-expanded AR molecules are more captured onto the DNA yet held less tightly. Lastly, ΔN AR displayed a high D bound that was ligand-insensitive, a minimal increase in F bound, and low rates of binding events that were also unchanged by ligand, indicating that, without the disordered NTD, the receptor retains only loose, shallow chromatin associations.

Next, we extracted the search time and track length parameters. Ligand-activated AR 24Q spent longer time in the mobile, chromatin-scanning state before committing to stable binding, reflecting an extended window of exploration (**Fig. 1k and 1l**). Conversely, both polyQ-expanded and ΔN AR showed reduced search times, likely consistent with premature exit from the search state and failure to adopt a search-competent state, respectively (**Fig. 1k and 1l**). Track lengths increased upon androgen exposure in both AR 24Q and AR 100Q, whereas they were unchanged in ΔN AR (**Fig. 1m**), further underscoring the NTD-dependence of ligand-driven chromatin capture.

Altogether, these findings support a model in which the AR NTD governs both the ligand-driven assembly of nuclear condensates and the productive search-and-capture dynamics of chromatin binding, with the polyQ expansion generating irregular, overly cohesive condensates that prematurely and aberrantly anchor AR in a loosely held, chromatin-associated state.

### Mapping AR chromatin binding and activity in skeletal muscle

We next proceeded to identify the genomic regions bound by AR upon androgen activation in human skeletal muscle cells. For this experiment, we developed a set of the hMyo allelic series whereby AR was tagged at the C terminal with a V5 tag (**Supplementary Fig. 1c and 1d**). We selected this epitope because in previous studies a single copy was sufficient to yield robust chromatin immunoprecipitation (ChIP) signal^36,37^. Having validated the assay specificity against a known AR binding site (**Supplementary Fig. 4a**), we performed ChIP sequencing (ChIP-seq) in proliferating myoblasts. This cell state provides uniform formaldehyde penetration, chromatin fragmentation and nuclear recovery, and critically avoids the confounding polyQ-dependent differentiation defects reported previously^38^.

A total of 44,907 high-confidence common binding regions were identified in hMyo expressing AR 24Q and AR 100Q after treatment with androgen (**Fig. 2a**). Notably, the binding intensity at transcriptional start sites (TSS) highlighted higher summit with comparable width of the average peaks in polyQ-expanded AR. This data further corroborates the markedly increased F bound SPT profiles of polyQ AR (**Fig. 1i**), overall representing more molecules captured at AR canonical sites. While the vast majority of binding sites were shared between wild type and polyQ-expanded AR, we also identified a number of differentially bound peaks (*P* < 0.05), which were clustered into two categories: only present in AR 24Q (*n* = 3,299, 6,76% of total peaks) or only present in AR 100Q (*n* = 595, 1,22% of total peaks) (**Fig. 2b; Supplementary Table 1**). Increased peak intensity was also evident within the 100Q-specific cluster, indicating that the gain in occupancy is not confined to sites shared with 24Q but extends genome-wide across polyQ-expanded AR binding (**Fig. 2b**). Upon ligand activation AR binding sites were predominantly located in the intronic (47,7%), intergenic (26,5%), and promoter regions (13,4%) (**Fig. 2c**), which is consistent with previous ChIP-seq experiments in prostate cancer setting^39,40^. Whereas the relative proportion of binding regions was virtually identical in the AR 24Q ‘only’ cluster, the AR 100Q ‘only’ cluster showed a shift towards distal intergenic regions (42.2%), primarily at the expense of promoter regions (6.4%) (**Fig. 2c**). To illustrate these categories at the locus level, representative genome-browser tracks are shown for each scenario (**Fig. 2d**).

**Fig. 2.**
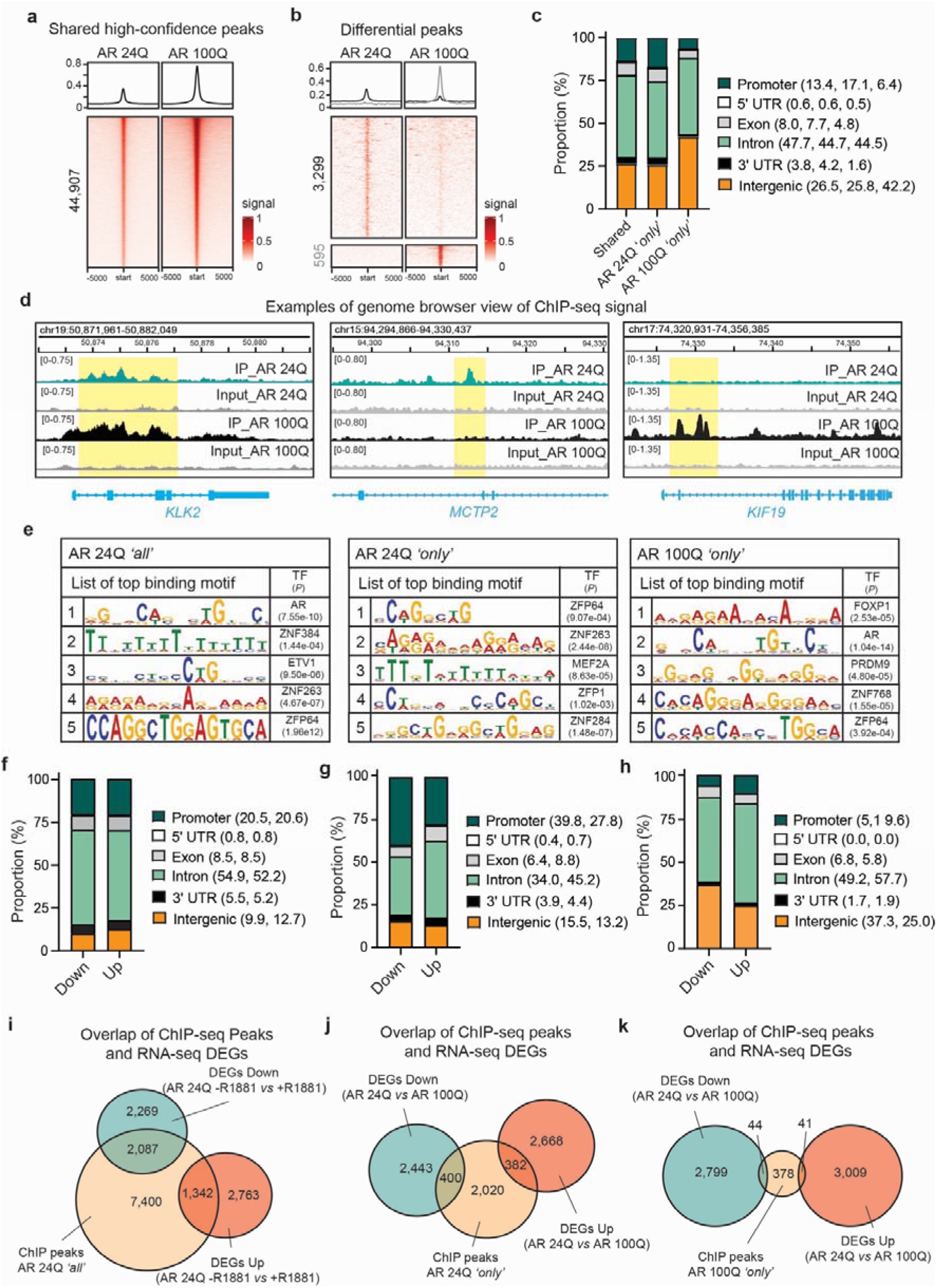
The AR cistrome and its coupling to the transcriptional output in human skeletal muscle cells. **a**, Average composite profiles (top) and heatmaps (bottom) of AR ChIP–seq signal at high-confidence binding regions shared between AR 24Q and AR 100Q hMyo after androgen treatment. Signal is centred on the peak summit and shown ± 5 kb; colour scale denotes normalised read density (0–1) (*n* = 2 biological replicates per genotype). **b**, Average composite profiles (top) and heatmaps (bottom) for the differentially bound regions: peaks called only in AR 24Q or only in AR 100Q. **c**, Genomic-feature distribution of the shared, AR 24Q-only and AR 100Q-only regions. Values in the legend give the proportion (%) for each feature in the order (shared, AR 24Q ‘*only*’, AR 100Q ‘*only*’). Intergenic peaks were assigned to the nearest transcription start site.**d**, Representative genome-browser views of AR ChIP–seq (IP) and input tracks for AR 24Q and AR 100Q at three loci illustrating each category: a shared peak at the androgen-responsive gene *KLK2* (left), an AR 24Q-only peak at *MCTP2* (middle) and an AR 100Q-only peak at *KIF19* (right). The peak region is highlighted in yellow; the track range is indicated in brackets. **e**, Top five *de novo* enriched sequence motifs for the shared (AR 24Q ’all’), AR 24Q-only and AR 100Q-only peak sets, with the matched transcription factor and enrichment *P* value. **f-h**, Genomic-feature distribution of AR peaks assigned to down-regulated *versus* up-regulated genes, for the shared AR 24Q ’all’ (**f**), AR 24Q-only (**g**) and AR 100Q-only (**h**) peak sets. Values in each legend give the proportion (%) per feature in the order (down, up). **i-k**, Integration of AR ChIP–seq binding with RNA-seq differential expression, shown as area-proportional Venn diagrams of the overlap between genes carrying an AR peak and differentially expressed genes (DEGs) (*n* = 3 biological replicates per genotype). **i**, Shared AR 24Q ’all’ peaks intersected with DEGs between untreated and androgen-treated AR 24Q (-R1881 *versus* +R1881). **j**, AR 24Q-only peaks intersected with DEGs between androgen-treated AR 24Q and AR 100Q. **k**, AR 100Q-only peaks intersected with the same AR 24Q versus AR 100Q DEG set. Numbers indicate gene counts in each compartment. *n* represents the number of experiments.

*De novo* motif analysis of the top shared AR-binding regions using the MEME suite^41^ revealed the AR responsive element sequence element as the highest score motif (*P* = 7.55e-10), confirming the specificity of the assay, together with ETS variant transcription factor 1 (ETV1) and GC-rich zinc-finger motifs (ZNF384, ZNF263, ZFP64), characteristic of the canonical AR cistrome and its established cofactors^42,43^ (**Fig. 2e**). Sites bound only by AR 24Q were enriched for Myocyte Enhancer Factor 2A (MEF2A) binding motif (*P* = 8.43e-5), a master regulator of skeletal-muscle homeostasis^44,45^ (**Fig. 2e**). Conversely, while retaining strong AR-motif enrichment polyQ expansion redistributes AR binding away from a MEF2-associated differentiation program, as previously described^46^, towards a distinct Forkhead box protein P1 (FOXP1)-associated circuit (*P* = 2.53e-05) (**Fig. 2e**).

To connect chromatin binding to transcriptional output, we integrated the AR ChIP-seq with RNA sequencing (RNA-seq) performed under the same conditions. Androgen treatment produced a transcriptional response with comparable numbers of down-regulated (*n* = 4,356) and up-regulated (*n* = 4,105) genes (**Supplementary Fig. 5a**). The same balanced split was seen when comparing androgen-treated AR 100Q *vs* AR 24Q (*n* = 2,843 down-regulated and *n* = 3,050 up-regulated genes; *P*adj < 0.01) (**Supplementary Fig. 5b**). When assessing the AR binding sites restricted to differentially expressed genes (DEGs), we observed an equal distribution across the various regions between the down- and up-regulated DEGs, with the vast majority occurring at intron and promoter regions (**Fig. 2f**). AR 24Q ‘only’ binding at DEGs was mainly enriched at promoters for down-regulated genes, with introns as the next most frequent category and a slight promoter-to-intron shift in up-regulated genes (**Fig. 2g**). AR 100Q ‘only’ binding, by contrast, was predominantly intronic and, secondarily, distal intergenic (peak assigned to its nearest transcription start site), consistent with a redistribution of the expanded AR away from promoter-proximal regulatory regions (**Fig. 2h**).

Because the relationship between binding and transcriptional output is most directly interpretable at promoters, we restricted the observations to high-confidence promoter AR peaks (binding window TSS ±1 kb). This analysis revealed that 67.7% promoter-bound genes showed no corresponding change in expression (*n* = 7,400), suggesting that most promoter binding is not accompanied by a productive transcriptional output (**Fig. 2i**). Upon androgen treatment, an AR promoter binding was present at 47.9% of down-regulated (*n* = 2,087) and 32.7% of up-regulated (*n* = 1,342) genes (**Fig. 2i**). Among genes differentially expressed between AR 100Q and AR 24Q, the sites bound only by AR 24Q coincided with a promoter peak at 14.1% of down-regulated (*n* = 400) and 12.5% of up-regulated (*n* = 382) genes, whereas the sites bound only by AR 100Q coincided with a promoter peak at just 1.5% of down-regulated (*n* = 44) and 1.3% of up-regulated (*n* = 41) genes (**Fig. 2j and 2k**). Thus, we concluded that the binding events at promoter regions unique to AR 100Q are frequently uncoupled from transcriptional output.

### AR mediates the effects of testosterone in muscle via AP-1

Because the anabolic effect of androgens on skeletal muscle requires a functional AR, we hypothesised that this outcome results from activation of a transcriptional programme driven by AR. Transcription-factor enrichment analysis (ChEA3)^56^ of the genes differentially expressed upon androgen treatment identified the activator protein 1 (AP-1) transcriptional complex as the most strongly enriched transcriptional regulator overall, as six of its components (FOSL2, JUND, JUN, FOS, FOSL1 and MAFK) were each significantly enriched (**Supplementary Fig. 7a**). The same AP-1 signature was recovered in an independent, large *in vivo* dataset comparing skeletal-muscle transcriptomes of bulls compared to castrated steers^47^ (**Supplementary Fig. 7b**). Gene Set Enrichment Analysis (GSEA)^48^ on the RNA-seq DEGs upon androgen treatment returned a significant positive enrichment of AP-1 genes (NES = 1.79, FDR *q*-value = 0.0039) (**Fig. 3a**).

**Fig. 3.**
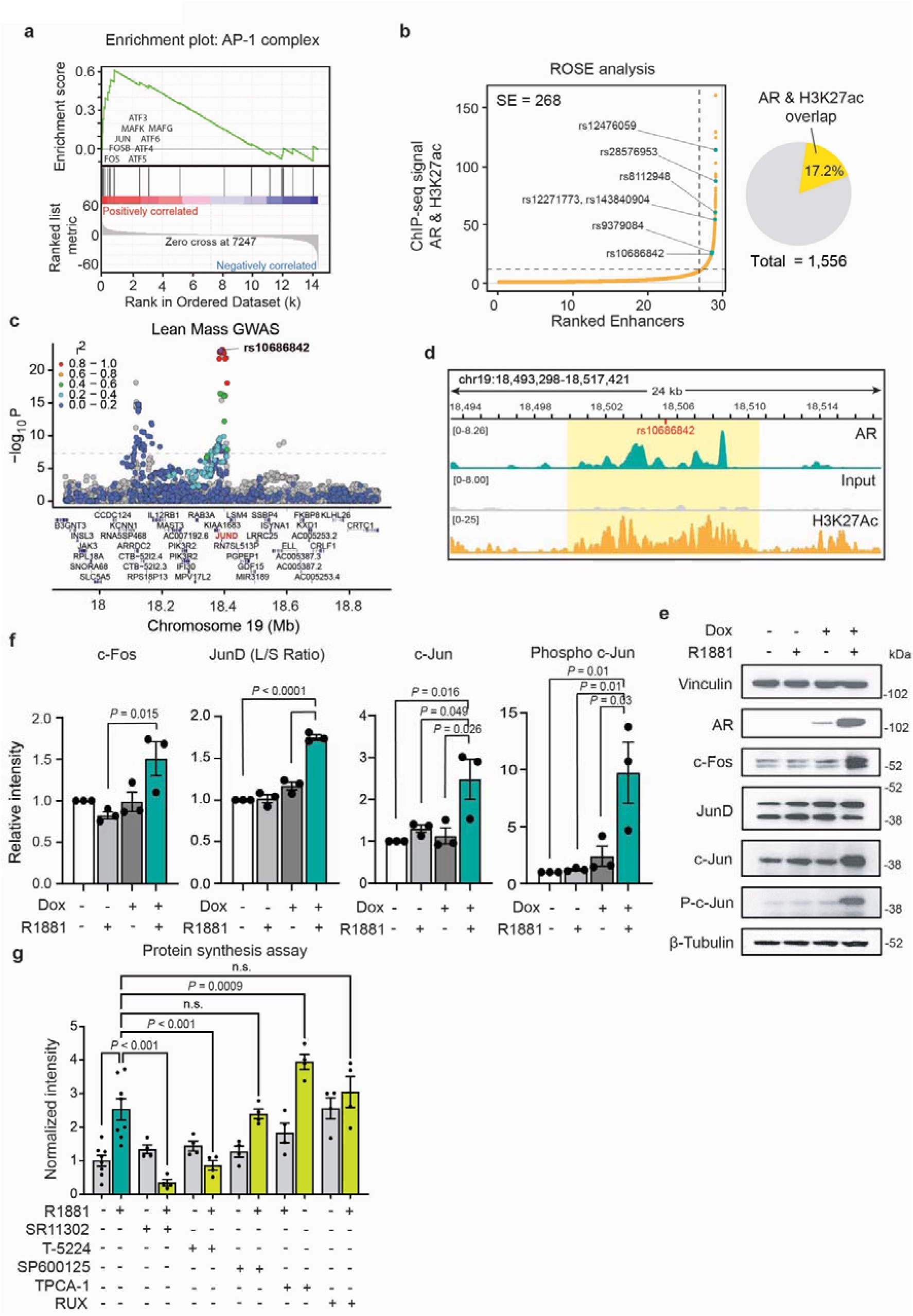
AR drives the androgen effects in skeletal muscle via AP-1. **a**, Gene-set enrichment analysis (GSEA) enrichment plot for the AP-1 complex gene set against genes ranked by differential expression upon androgen treatment. The running enrichment score (top), gene positions (middle) and ranked-list metric with the zero-crossing at rank 7,247 (bottom) are shown; leading-edge AP-1 members (*FOS, ATF3, FOSB, MAFK, MAFG, JUN, ATF4, ATF5, ATF6*) are indicated. **b**, Ranking of super enhancers (SE) by the ROSE algorithm, combining H3K27ac and AR ChIP–seq signal in androgen-treated AR 24Q hMyo; SE lie above the inflection point (dashed line). Trait-associated SNPs from the lean-mass GWAS^49^ that fall within these regions are labelled. The pie chart shows percentage of AR– H3K27ac co-bound regions with super-enhancer features. **c**, Regional association plot for the lean-mass GWAS across the chromosome 19 locus, plotting −log₁₀(*P*) against genomic position. The lead variant rs10686842 is highlighted; other SNPs are coloured by linkage disequilibrium (r²) with the lead SNP (legend, top left). Genes in the interval are shown below; *JUND* is highlighted in red. **d**, Genome-browser view of AR (IP), input and H3K27ac ChIP–seq tracks across the *JUND* locus (chr19:18,493,298-18,517,421). The region containing rs10686842 (red mark) is highlighted in yellow. **e**, **f**, Representative immunoblots of AR, c-Fos, JunD, c-Jun and phospho-c-Jun in AR 24Q hMyo as a function of doxycycline-induced AR expression (Dox) and androgen (R1881). Vinculin and β-tubulin serve as loading controls; molecular-weight markers (kDa) are shown at right. **f**, Quantification of the immunoblots in e for c-Fos, the JunD long/short (L/S) isoform ratio, c-Jun and phospho-c-Jun, expressed as relative intensity. Bars show mean ± s.e.m. with individual biological replicates overlaid (*n* = 3); *P* values are from one-way ANOVA with Tukey’s multiple-comparisons test. **g**, Nascent protein synthesis measured by the Click-iT^TM^ Plus OPP Alexa Fluor^TM^ assay (Thermo Fisher) in AR 24Q hMyo, with and without R1881, under DMSO or the inhibitors SR11302 and T-5224 (AP-1), SP600125 (JNK), TPCA-1 (IKK/NF-κB) and ruxolitinib (RUX; JAK1/2), each at 1 µM. The same results were observed at 100 nM (not shown). Bars show mean ± s.e.m. of fluorescence intensity normalised to -R1881, DMSO-treated, with individual biological replicates overlaid (*n* = 3 – 8). *P* values are from one-way ANOVA with Tukey’s multiple-comparisons test; n.s., not significant.

In parallel, to prioritise functionally relevant sites in an unbiased manner, we intersected the AR cistrome with genetic determinants of muscle mass and strength in humans. From the largest-to-date genome-wide association study (GWAS), resulting in a list of 1,059 single nucleotide polymorphisms (SNPs) across 799 loci significantly associated with lean muscle mass in 450,243 UK Biobank participants^49^, we extracted the common variants mapping onto our high-confidence AR binding regions (*n* = 12).

Overlapping of our AR binding sites with H3K27ac peaks, marker of active regulatory activity^50^, from a previously published dataset obtained in the psoas muscle of a male adult individual^51^, we found that most of these trait-associated SNPs (*n* = 7/12) lay within regions bearing super enhancer features, as defined by the Rank Ordering of Super-Enhancers (ROSE) pipeline (v0.1; https://hpc.nih.gov/apps/ROSE.html). Super enhancers (SE) are large clusters of high transcription factor occupancy enriched for disease- and trait-associated variants and functioning as master controllers of cell identity and disease state^52,53^ (**Fig. 3b**).

Using the Functional Mapping and Annotation of GWAS (FUMA) platform^54^ and Multi-marker Analysis of GenoMic Annotation (MAGMA) statistical tool for gene-based association analysis^55^, we assigned one of the mapped candidate variants, rs10686842, to *JUND*, which encodes the JunD protein, a core component of the AP-1 complex (**Fig. 3c and 3d**). Open Targets L2G ranked *JUND* as the most likely causal gene at the rs10686842 locus^56^ (L2G = 0.397 vs 0.306 for the next candidate, *IQCN*). The score was driven by TSS proximity together with colocalising expression- and splicing-QTL (eQTL) evidence for *JUND*, whereas the negligible variant-effect-predictor contribution indicates a non-coding, regulatory mechanism consistent with AR action through the associated super-enhancer (**Supplementary Fig. 6a**). Importantly, *JUND* mRNA levels were increased upon androgen treatment, suggesting true regulation (**Supplementary Fig. 6b**). Thus, both TF activity inference from AR-dependent gene expression and human-genetic prioritisation of a muscle-mass locus converged on the same AP-1 complex node.

We next sought to assess whether the AP-1 complex is activated by testosterone. Androgen treated hMyo cells showed a significant increase in c-Fos, c-Jun and phospho-c-Jun, and an increased ratio of the long JunD isoform relative to the short isoform, which is the more potent transcriptional activator^57^ (**Fig. 3e and 3f**; **Supplementary Fig. 8a**). Critically, this induction occurred only in the presence of ligand-activated AR, indicating strict dependence on AR transcription.

Lastly, we tested whether this programme is absolutely required for mediating the androgen-driven anabolic response in skeletal muscle, applying a panel of pathway inhibitors. Reasoning that testosterone increases muscle mass by promoting protein translation, we observed that exposure to androgen increased the nascent protein synthesis rate (**Fig. 3g**). This elevation was abolished by two mechanistically distinct AP-1 inhibitors: T-5224, which blocks AP-1 DNA binding^58^, and SR11302, which blocks AP-1 transactivation^59^, establishing a specific requirement for AP-1 transcriptional activity (**Fig. 3g**). In contrast, inhibition of the Nuclear Factor kappa B (NF-κB) pathway with the IκB kinase beta (IKKβ) inhibitor TPCA-1^60^ further increased protein synthesis, indicating that NF-κB acts as an opposing, catabolic arm as previously shown^61^. Conversely, the c-Jun N-terminal kinase (JNK) inhibitor SP600125^62^ and the Janus kinase 1/2 (JAK1/2) inhibitor ruxolitinib^63^ had no effect, indicating that AP-1 activity is not rate-limited by JNK-dependent c-Jun phosphorylation and does not require JAK–STAT signalling (**Fig. 3g**). Together, these data place AP-1 transcriptional activity as the required downstream effector of ligand-activated AR, counterbalanced by NF-κB, in driving the anabolic response.

### Polyglutamine-expanded AR drives skeletal muscle atrophy through aberrant EDA2R transcription

Having established that AR chromatin binding is a key step mediating the effects of testosterone in skeletal muscle, we next examined the role of the polyQ tract. To do so at population scale, we leveraged a sub-group of individuals from the UK Biobank imaging cohort^64–67^, where body composition was quantified by fully automated volumetric analysis, as previously described^68^, while liver MRI data were analysed with LiverMultiScan® (Perspectum, Oxford, UK) (*n* = 29,792 males and 34,099 females).

We assessed associations between *AR* CAG-repeat length, determined using ExpansionHunter^69^, and imaging-derived phenotypes by multivariable linear regression, modelling repeat length as a continuous predictor so that each coefficient represents the expected change in the phenotype per additional CAG repeat. Models were adjusted for age and circulating testosterone to control for their confounding influence on skeletal muscle. Notably, in males, CAG-repeat length was itself positively associated with testosterone levels, most likely by a compensatory response to reduced receptor efficiency (ρ = 0.07, 95% CI = 0.06 - 0.08, *n* = 21,059, *P* = 4.19e25).

Because androgen signalling differs substantially between the sexes, we fitted separate models in men and women for each clinical and imaging phenotype. In males, muscle fat infiltration (MFI) increased significantly with each additional CAG repeat, whereas no such association was detected in females (**Fig. 4a**). Body mass index (BMI) and liver proton-density fat fraction (PDFF) likewise rose with increasing CAG-repeat length, indicating a male-specific effect of repeat size on fat distribution and organismal metabolism (**Fig. 4a**). Interestingly, these human genetic associations mirror the alterations reported in SBMA, in which skeletal muscle displays disrupted energy metabolism^70,71^, quantitative muscle MRI shows progressive intramuscular fat infiltration^72,73^, and the liver develops non-alcoholic fatty infiltration^74,75^.

**Fig. 4.**
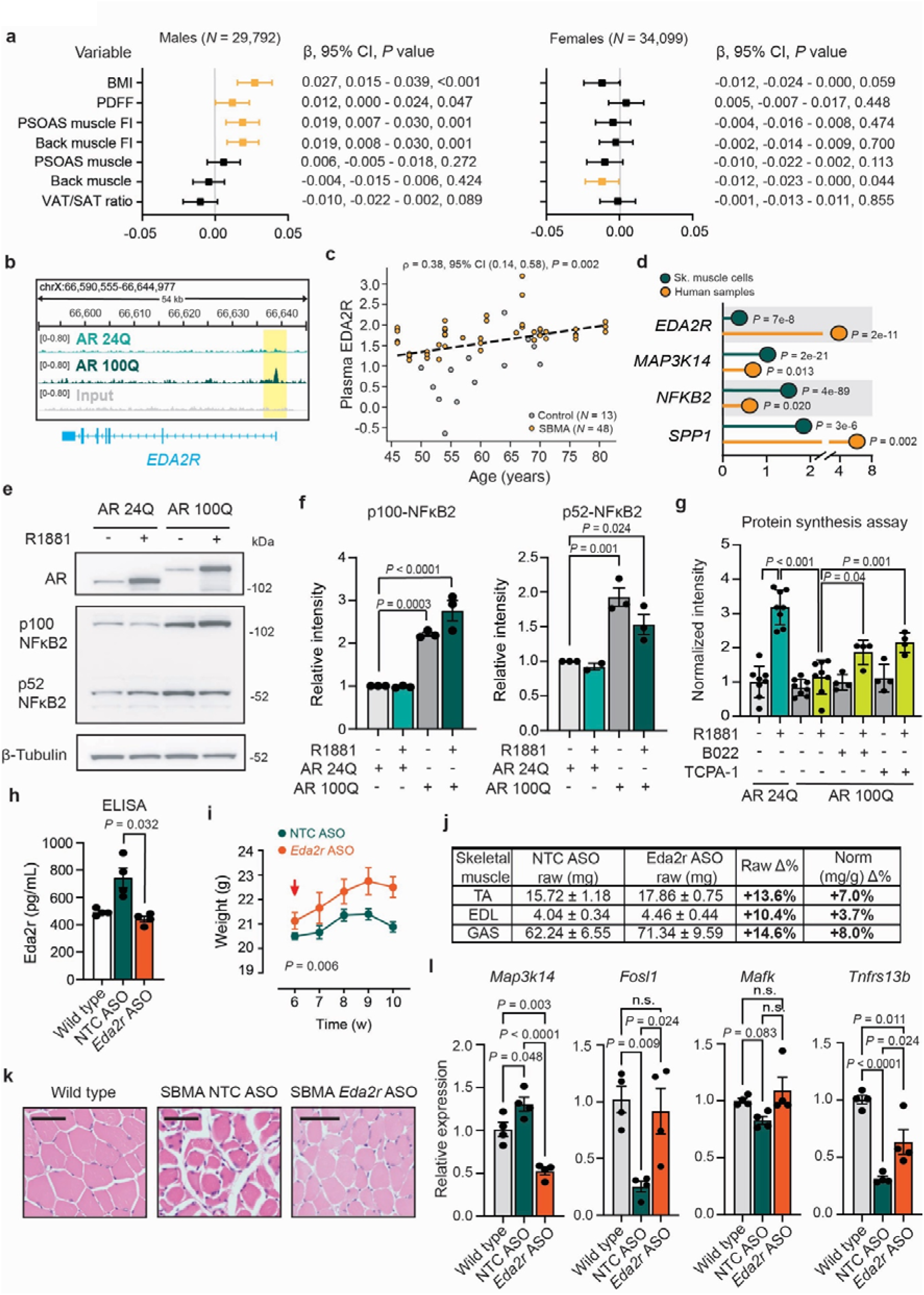
Transcriptional activation of *EDA2R* drives muscle atrophy in SBMA. **a**, Sex-stratified associations between the *AR* CAG-repeat length and body-composition traits in a cohort of individuals from the UK Biobank. Forest plots show the per-allele effect size (β) with 95% confidence interval; β, 95% CI and P value are listed at right for each trait. Coloured symbols denote nominally significant associations (*P* < 0.05). BMI: body mass index, PDFF: proton density fat fraction, MFI: muscle fat infiltration, VAT: visceral adipose tissue, SAT: subcutaneous adipose tissue. **b**, Genome-browser view of AR (IP, AR 24Q and AR 100Q) and input ChIP-seq signal at the *EDA2R* locus (chrX:66,590,555-66,644,977); the AR 100Q-selective peak is highlighted in yellow. **c**, Plasma EDA2R protein (relative units) versus age in SBMA patients (*N* = 48) and controls (*N* = 13); each point is one sample. The dashed line is the linear fit; Spearman ρ = 0.38 (95% CI 0.14–0.58), *P* = 0.002. **d**, Log₂ fold change in expression of *EDA2R*, *MAP3K14*, *NFKB2* and *SPP1* from RNA-seq, in polyQ-expanded versus wild-type AR skeletal-muscle cells (Sk. muscle cells, teal) and in SBMA versus control human samples (orange), shown as a lollipop plot with the corresponding *P* value for each gene and dataset (DESeq2 Wald test, adjusted *P*). **e**, Representative immunoblots of AR, p100-NF-κB2 and its processed p52-NF-κB2 product in AR 24Q and AR 100Q hMyo, with and without androgen. β-tubulin is the loading control; molecular-weight markers (kDa) are shown at right. **f**, Quantification of **e** for p100-NF-κB2 (left) and p52-NF-κB2 (right), expressed as relative intensity. Bars show mean ± s.e.m. with individual biological replicates overlaid (*n* = 3); *P* values from one-way ANOVA with Tukey’s multiple-comparisons test. **g**, Nascent protein synthesis in AR 24Q and AR 100Q hMyo ± R1881, treated with the NIK inhibitor B022 or the IKKβ inhibitor TPCA-1 (each at 100 nM) or DMSO vehicle. Bars show mean ± s.e.m. of normalised OPP fluorescence intensity with individual replicates overlaid (*n* = 4–8); *P* values from one-way ANOVA with Tukey’s multiple-comparisons test. **h**, Plasma EDA2R measured by ELISA (pg ml⁻¹) in wild-type mice and SBMA mice treated with non-targeting control (NTC) ASO or *Eda2r*-targeting ASO. Bars show mean ± s.e.m. (*n* = 4–5 mice per group); *P* value from one-way ANOVA. **i**, Body weight over weeks 6-10 in SBMA mice receiving NTC ASO (teal) or *Eda2r* ASO (orange); the red arrow marks the start of ASO dosing. Data are mean ± s.e.m. (*n* = 4–5 per group); *P* = 0.006 two-way repeated-measures ANOVA. **j**, Wet weights of hindlimb skeletal muscles (tibialis anterior, TA; extensor digitorum longus, EDL; gastrocnemius, GAS) from SBMA mice treated with NTC ASO or *Eda2r*-targeting ASO (n = 5 per group). Raw values are mean ± s.e.m. absolute wet weight (mg); Δ% is the percentage difference in the *Eda2r* ASO group relative to NTC ASO. Because body weight differed between groups (two-tailed unpaired t-test, *P* = 0.022), muscle weights were normalized to body weight; Norm Δ% is the percentage difference in the body-weight-normalized weight (mg g⁻¹). **k**, Representative haematoxylin and eosin–stained cross-sections of TA muscle from wild-type, SBMA NTC ASO and SBMA *Eda2r* ASO mice, showing rescue of myofibre morphology by *Eda2r* silencing. Scale bar, 50 µm. **l**, RT–qPCR of mouse *Ma3k14, Tnfrs13b*, *Fosl1,* and *Mafk* in muscle from wild-type and SBMA mice treared with NTC ASO or *Eda2r* ASO, as relative expression. Bars show mean ± s.e.m. (*n* = 4-5); *P* values from one-way ANOVA with Tukey’s multiple-comparisons test; n.s., not significant.

We next sought to dissect the consequences of a pathologically expanded polyQ tract in skeletal muscle. To this end, we adopted the following logic: First, because muscle atrophy occurs in SBMA but not in conditions of AR insensitivity, we inferred a gain-of-function mechanism and therefore focused on the sites differentially bound by mutant AR (**Fig. 2a-c**). Second, we intersected these regions of differential or new binding upon polyQ-expanded AR with transcriptomic data from the human muscle cells, the SBMA mouse model, and patient muscle biopsies (**Supplementary Fig. 9a**). Across all datasets, the most strongly differentially expressed target was ectodysplasin A2 receptor (*EDA2R*), driven by aberrant AR binding at its promoter (**Fig. 4b**). Notably, EDA2R up-regulation is associated with dysregulated metabolism as hallmarks of aging^76,77^ and has been recently shown to promote muscle atrophy in tumour-bearing mice and in patients with cancer cachexia via activation of the non-canonical NF-κB pathway^78^.

Consistently, a recent plasma proteomic study of SBMA patients reported that EDA2R protein was significantly elevated relative to age- and sex-matched controls and correlated inversely with clinical severity, as measured by the SBMA functional rating scale^79^. In this cohort, plasma EDA2R levels also rose with age and were consistently higher in SBMA patients than in matched controls (**Fig. 4c**).

We confirmed that EDA2R up-regulation activated the non-canonical NF-κB pathway, verifying induction of pathway nodes (*MAP3K14, NFKB2* and *SPP1*) in both the human skeletal muscle cells and SBMA muscle biopsies relative to age- and sex-matched controls (**Fig. 4d**). Non-canonical NF-κB activation depends on NF-κB-inducing kinase (NIK), which drives processing of p100–NFκB2 into the active p52–NFκB2 form and nuclear translocation of the p52–NFκB2–RelB complex^80^. Accordingly, polyQ AR produced marked activation of NF-κB signalling, evidenced by increased expression and NIK-driven processing of p100– NFκB2 to p52–NFκB2 (**Fig. 4e and 4f**). Notably, EDA2R ligand engagement is also known to activate the JNK/c-Jun N-terminal kinase cascade, converging on the AP-1 complex^78,81,82^. In line with previous evidence in SBMA and other polyQ disorders^83,84^, we observed further increased c-Fos/Jun activation in SBMA muscle (**Supplementary Fig. 10a**). Finally, androgen treatment reduced the overall rate of protein synthesis in SBMA hMyo (**Fig. 4g**), an effect partially restored by NF-kB inhibitors B022^85^ and TCPA-1^60^, confirming its dependence on EDA2R-driven signalling (**Fig. 4g**). Together, these results point to a gain-of-function mechanism of muscle atrophy in SBMA, driven by aberrant transcriptional activation of the NF-κB pathway.

To test whether EDA2R-mediated NF-κB activation is sufficient to drive the SBMA muscle-atrophy programme, we asked whether silencing *Eda2r* could reverse muscle atrophy in the SBMA transgenic mice carrying an expanded AR (AR 100Q), a model which faithfully recapitulates key disease features, including testosterone-driven pathogenesis and progressive muscle atrophy and weakness^86^. We first confirmed that *Eda2r* mRNA was elevated by approximately four-fold in this model, reinforcing the consistency of this finding across disease models and human patients (**Supplementary Fig. 11a**). Using OligoAI, an artificial intelligence model we recently developed to predict antisense oligonucleotide (ASO) *in vitro* knockdown efficacy^87^, we selected the highest-scoring gapmer ASOs targeting *Eda2r* (**Supplementary Fig. 12a**). We confirmed knock-down activity in cells (**Supplementary Fig. 12b**) and advanced the single ASO active against both human and murine *EDA2R* into the SBMA mouse model (**Supplementary Fig. 12a and 12c**). Peripheral administration of the *Eda2r*-targeting *versus* non-targeting ASO (30 mg/kg, twice weekly) from disease onset at 6 weeks of age to 10 weeks in SBMA male mice (*n* = 4-5 per arm), normalized serum *Eda2r* levels measured by ELISA compared to wild type age- and sex-matched littermates (**Fig. 4h**). Treatment also significantly attenuated the weight loss characteristic of this model (**Fig. 4i**), with improvements in wet weights of key skeletal muscles (both raw and normalised to overall body weight) (**Fig. 4j**) and histology architecture (**Fig. 4k)**. Finally, mRNA levels of downstream effectors of both the NF-κB (*Map3k14, Tnfrs13b*) and AP-1 (*Fosl1, Mafk*) pathways were restored to normal, confirming on-target engagement (**Fig. 4l**). Collectively, these data demonstrate that pathologically expanded polyQ AR acts through a toxic gain-of-function mechanism, driving a shift toward altered metabolism and fat deposition and transcriptionally triggering a muscle-atrophy programme via the non-canonical NF-κB pathway.

## Discussion

AR mediates the potent anabolic effects of androgens in skeletal muscle, yet the molecular underpinnings remain undefined. By combining super-resolution imaging, single-molecule tracking, genome-wide binding and expression maps, and human population genetics, we trace a continuous chain of causation from receptor biophysics to muscle phenotype. Ligand-activated AR assembles nuclear condensates through its intrinsically disordered NTD, converts chromatin scanning into stable, productive capture, and channels its output through the AP-1 complex to drive muscle growth. AP-1 is a dimeric complex assembled from Jun, Fos, ATF and MAF-family subunits whose composition dictates its output, and it functions as a master integrator of the extracellular signals governing proliferation, differentiation, survival and death^88,89^.

While an intact NTD is essential for correct condensate formation and chromatin scanning, a pathologically expanded polyQ tract switches the receptor’s anabolic programme into a catabolic one, offering an ideal model of study of AR function. Although historically classified as a lower-motor-neuron disease, converging evidence has indeed established skeletal muscle as a critical site of toxicity in SBMA: muscle-restricted expression of polyQ AR recapitulates systemic and even motor neuron disease features^90^, peripheral suppression of AR in muscle rescues the phenotype across mouse models^71,91^, and muscle-directed IGF-1 is protective^92,93^.

Our data support a model by which polyQ expanded AR is captured more frequently but less tightly onto the chromatin, representing the kinetic signature of sequestration into viscous, over-cohesive condensates. Importantly, such increased chromatin occupancy by mutant AR does not translate into a single directional shift in gene expression, with a large fraction of binding events with no detectable transcriptional consequence. This is the behaviour expected of a *bona fide* transcription factor, whose effect at any given locus depends on local co-factor availability, chromatin state and enhancer–promoter architecture rather than on occupancy alone, resulting in a genome-wide rewiring of the AR regulatory logic, rather than a simple gain or loss of activity. This complexity aligns AR with the emerging principle that repeat expansions in transcription factor IDRs act by altering condensate material properties rather than by a simple mechanism of loss of function^19^. Notably, we also observed that a larger proportion of expanded AR molecules fails to translocate into the nucleus, increasing the cytoplasmic burden of aggregation-prone polyQ AR, offering a mechanistic link to the heightened cytoplasmic proteostatic and stress responses previously reported in SBMA^94–99^.

A distinctive feature of this rewiring is a spatial redistribution of expanded AR away from promoters and towards distal intergenic sites, most of which correspond to enhancer regions. Unlike promoters, these regulatory regions are built predominantly from low-affinity, degenerate binding motifs, and this suboptimal syntax is key to enable their tissue-specific, signal-responsive and combinatorially controlled activity^100,101^. The corollary is that enhancer regions are intrinsically more permissive and promiscuous binding platforms than the constrained, high-affinity core motifs of promoters. We therefore hypothesise that a TF whose search-and-capture behaviour has been altered by the repeat expansion (i.e.: capturing DNA more frequently but binding less specifically) is therefore expected to drift into these low-stringency distal compartments, where transient, degenerate engagement is tolerated. The observed promoter-to-enhancer redistribution is thus a consequence of the altered condensate and binding kinetics measured directly and may help explain why so many of the new binding events are transcriptionally unproductive.

The mechanistic evidence assembled to date on the molecular mechanisms of muscle atrophy in SBMA has pointed almost exclusively towards a loss of function, with recent studies showing defective excitation-contraction coupling and mitochondrial respiration upon polyQ AR^102^ and reduced transcriptional activation of the myogenic master regulator MEF2^46^, which was also confirmed by our genome-wide analysis.

However, a purely loss-of-function model cannot account for the fact that AR deficiency itself does not cause muscle atrophy. Here we identify aberrant promoter binding of expanded AR, resulting in *EDA2R* up-regulation which is consistent across cellular models, SBMA transgenic mice, and patient muscle biopsies. ASO-mediated silencing of *Eda2r* normalised NF-κB effectors and attenuated muscle wasting in a proof-of-principle study in SBMA mice, suggesting that this mechanism is causal and actionable. Importantly, EDA2R has been recently shown to promote muscle atrophy in cancer cachexia by activating the non-canonical NF-κB axis^78,80^, while in parallel activating the JNK cascade, the principal upstream kinase driving c-Jun phosphorylation^81^, placing SBMA on a shared common pathway of muscle wasting. Our data position this AP-1 arm as the required transducer of the anabolic AR signal, counterbalanced by the catabolic NF-κB arm^61^, framing muscle mass as a transcriptionally set balance between opposing programmes that AR can tip towards growth or wasting depending on the size of the repeat expansion in the NTD.

Beyond its role in skeletal muscle homeostasis, EDA2R is rapidly emerging as a marker of cellular senescence, inflammation and aging^76^. Interestingly, in a recent plasma proteomic profiling in SBMA individuals relative to age- and sex-matched unaffected controls, EDA2R was found markedly elevated and correlate with clinical scales^79^. Its direct engagement by expanded AR therefore implicates accelerated, transcriptionally-driven aging of muscle as a driver of SBMA, consistent with the adult-onset, progressive nature of the disease and with the systemic metabolic and ectopic-fat signature observed in patients^70,72,74^ and point towards circulating EDA2R as a promising biomarker.

In our study we have deliberately focused on the transcriptional activity of AR, its condensate behaviour, cistrome and downstream gene programmes, yet AR also exerts rapid, non-genomic actions through various signalling cascades^103–105^, which are not addressed here; how they are affected by the repeat expansion remains an open and important question. In addition, our cellular work was carried out in proliferating myoblasts: While this choice ensured uniform chromatin accessibility and avoided the confounding effects of polyQ dependent differentiation defects, it does not capture the additional layer of regulatory complexity introduced by myogenic differentiation. Our work opens several follow-up questions, such as: Which are the functional components of AR condensates? Can condensate-targeting therapies enhance or restore AR activity? What is the composition of the AP-1 complex upon androgen stimulation and how does this change in the context of SBMA? How does AR genomic occupancy differ across tissues?

By linking IDR repeats to condensate dynamics, cistrome rewiring and a defined atrophy pathway, this work not only provides a molecular understanding of the anabolic function of AR in skeletal muscle but also sheds light on a fundamental mechanism of disease pathogenesis in SBMA and offers a broader framework to study transcriptional activity of disease genes harbouring microsatellite repeats.

## Materials and Methods

### Cell Culture

Immortalized human myoblasts (MRC CNMD Biobank London, L954/1284 M-I) were maintained in skeletal muscle cell growth medium (PromoCell) supplemented with supplement mix (PromoCell), 10% (v/v) fetal bovine serum (FBS; Gibco), 1× GlutaMAX (Gibco), and 1× antibiotic antimycotic (Gibco). For experimental conditions, myoblasts were exposed to 1 µg/mL doxycycline (Sigma-Aldrich) and 10 nM R1881 (Sigma-Aldrich) or vehicle control in FBS-free growth media for 24 or 48 hours as indicated.

### Plasmid Construction

To generate doxycycline-inducible AR expression constructs, cDNAs encoding AR 24Q, AR 66Q, AR 100Q, or an AR variant lacking the N-terminal domain (ΔN) were cloned by restriction enzyme-based cloning or Gibson assembly into a PiggyBac backbone, under the control of a tetracycline response element (TRE) and flanked by PiggyBac terminal repeat sequences. Constructs were generated with either a HaloTag or V5 epitope tag. All plasmid constructs were sequence verified by whole-plasmid Nanopore30 sequencing (Source Genomics, Cambridge, UK).

### Cell Line Generation

AR knock-out myoblast lines were generated using ribonucleoprotein (RNP) complexes. Single-guide RNA (sgRNA) targeting the endogenous androgen receptor (Integrated DNA Technologies; **Supplemental Table 1**) was combined with AltR S.p. Cas9-GFP V3 (Integrated DNA Technologies) at a gRNA:Cas9 ratio of 1.2:1 in RNAiMax (Thermo Scientific). Wild-type myoblasts were transfected with RNPs at 10nM final concentration in Opti-MEM (Gibco). GFP-positive cells were isolated using the Sony MA900 Cell Sorter. Prior to sorting, cells were harvested and resuspended in culture media at 1x10 cells/mL. The sorter was calibrated using setup beads (BD™ CompBeads Anti-Mouse Ig, κ/Negative Control Compensation Particles Set) with appropriate fluorescence compensation. GFP fluorescence was detected using a 488nm laser. Gates excluded debris and doublets based on forward and side scatter parameters. Sorted cells underwent single-cell seeding via serial dilution, and expanded clones were assessed for endogenous AR expression relative to wild-type myoblasts by Western Blot.

Stable human myoblast lines expressing V5- or HT-tagged AR 24Q, AR 66Q, AR 100Q, or AR ΔN were generated using the PiggyBac transposon system. Human myoblasts were transfected at approximately 20–30% confluence using Lipofectamine™ 2000 (Thermo Fisher Scientific). Cells were co-transfected with 1000ng of the respective AR expression plasmid and 100ng of the PB210 PiggyBac transposase plasmid (PB210PA-1; Stratech Scientific). Following transfection, cells were subjected to selection with 10 µg/mL puromycin. Puromycin-resistant cells were subsequently single-cell cloned, and individual monoclonal lines were expanded. AR transgene expression was induced with 1 µg/mL doxycycline and confirmed by Western blotting.

### Androgen receptor luciferase reporter assay

To assess AR-dependent transcriptional activity, HEK293T cells were transiently transfected using Lipofectamine™ 2000 (Thermo Fisher Scientific) in Opti-MEM™ I Reduced Serum Medium (Gibco) with 0.5 µg of an androgen response element (ARE)-driven firefly luciferase reporter plasmid, 0.5 µg of pRL-TK Renilla luciferase control plasmid (Promega), and 0.5µg of AR 24Q-HT or AR 100Q-HT expression plasmid. Following 24 hours of transfection, cells were treated with 1 µg/mL doxycycline and, where indicated, 10 nM R1881 and/or 0.5 nM leptomycin B (LMB) for 24 hours. Firefly and Renilla luciferase activities were subsequently measured using the Dual-Glo® Luciferase Assay System (Promega) according to the manufacturer’s instructions. Luciferase activity was measured using a microplate spectrophotometer (CLARIOstar, BMG LabTech). Renilla luciferase activity was used as an internal normalization control, and AR-dependent transcriptional activity was expressed as firefly luciferase activity normalized to the corresponding Renilla luciferase signal.

### Fluorescence recovery after photobleaching (FRAP)

Fluorescence recovery after photobleaching (FRAP) was performed on human myoblasts stably expressing AR 24Q-HT, AR 66Q-HT, AR 100Q-HT or AR ΔN-HT. Cells were treated with 1 µg/mL doxycycline and 10 nM R1881 or vehicle in FBS-free growth media for 24 hours. AR-HT constructs were labelled with JF549 HT ligand (Promega) for live-cell fluorescence imaging. Cells were imaged using a ZEISS Axio Observer.Z1/7 laser-scanning confocal microscope equipped with a C-Apochromat 40×/1.20 NA water-immersion objective. Ten pre-bleach frames were acquired before a 1µm-diameter circular region of interest (ROI) was photobleached using the 561-nm laser at 25% power for 10 iterations. Fluorescence recovery within the bleached ROI was subsequently monitored by time-lapse imaging for 15 seconds. Fluorescence intensity within the bleached ROI was quantified over time and normalized to the pre-bleach fluorescence intensity. Normalized recovery curves were used to determine fluorescence recovery kinetics.

### Τau-STED imaging and image analysis

Tau-STED Microscopy was performed on human myoblasts stably expressing AR 24Q-HT, AR 66Q-HT, AR 100Q-HT or AR ΔN-HT. Cells were treated with 1µg/mL doxycycline and 10nM R1881 or vehicle in FBS-free growth media for 24 hours. AT-HT constructs were labelled with JF549 HT ligand (Promega) for live-cell imaging.

Super resolution imaging was performed using a Leica TCS SP8 STED microscope (Leica microsystems, Mannheim Germany) with a FALCON (FAst Lifetime CONtrast) module and an HC PL APO 86x/1.20 NA water-immersion objective with a motorised correction collar. Fluorescence was excited using a pulsed (80 MHz) supercontinuum 561-nm white-light laser (WLL) and depleted using a 660-nm STED laser. Tau-STED images were then generated using fluorescence lifetime and subsequently analysed using the Huygens Object Analyzer (Scientific Volume Imaging, The Netherlands). Individual fluorescent objects were identified and separated using watershed segmentation. For each segmented object, voxel number, voxel area, integrated and mean fluorescence intensity, centre-of-mass coordinates, length, width, and morphological parameters describing object roughness and roundness were extracted for quantitative analysis.

### Single-molecule imaging and tracking

Single-molecule imaging, tracking, diffusion analysis, and estimation of diffusion state residence times of HaloTag-AR were performed as previously described^104^. Briefly, AR-HT constructs were labelled with JF549 and imaged by highly inclined illumination microscopy using a Zeiss Axio Observer Z1 equipped with a 100×/1.46 NA objective and an iLas² illumination system (Cairn Optics). JF549 was excited using a 561-nm laser, and fluorescence emission was collected at 580–620 nm using an Andor iXon+ EMCCD camera. Time-series images were acquired at 10 Hz for 800 frames. Single AR-HT molecules were detected and tracked using Bayesian segmentation and proximity-based track linking, with tracks outside the nucleus or out of focus excluded from subsequent analyses. Diffusion kinetics were determined from 10 Hz datasets using Spot-On and fitted to a two-state model to estimate the chromatin-bound and freely diffusing fractions and their respective diffusion coefficients. A two-state hidden Markov model (HMM) was used to classify track segments as bound or freely diffusing. Distributions of free diffusion segment durations were corrected for photobleaching and used to estimate the characteristic search time. Binding events, defined as transitions from the diffusing to bound state, were identified in the segmented tracks and used to calculate the binding rate per area for each experimental condition. Parameter variability was estimated by bootstrap resampling of tracks (1,000 resamples per experimental condition).

### RNA extraction, cDNA synthesis, and RT-qPCR

For skeletal muscle samples, 10mg of tissue was homogenized in 1-thioglycerol homogenization solution using a Precellys tissue homogenizer (Bertin Instruments). Total RNA was isolated from homogenized muscle tissue or cultured cells using the Maxwell® RSC simplyRNA Tissue Kit on a Maxwell® RSC Instrument (Promega), according to the manufacturer’s instructions. For cDNA synthesis, 1 µg of total RNA was reverse-transcribed using the High-Capacity cDNA Reverse Transcription Kit (Applied Biosystems) according to the manufacturer’s instructions. RT-qPCR was performed using Power SYBR™ Green PCR Master Mix (Applied Biosystems) on a QuantStudio™ Real-Time PCR System (Applied Biosystems) according to the manufacturer’s recommended cycling conditions. Gene-specific primer sequences are provided in **Supplementary Table 2**.

### Immunoblotting

Whole-cell lysates were prepared in RIPA buffer (Thermo Fisher Scientific) supplemented with cOmplete™ EDTA-free Protease Inhibitor Cocktail and PhosSTOP™ (Roche) phosphatase inhibitors. Protein concentrations were determined using the Pierce™ BCA Protein Assay Kit (Thermo Fisher Scientific). Equal amounts of protein (10µg) were resolved by SDS-PAGE using 4–12% NuPAGE™ gels (Thermo Fisher Scientific) and electrotransferred onto nitrocellulose membranes using the Novex™ XCell SureLock™ Mini-Cell and XCell II™ Blot Module (Invitrogen). Membranes were blocked for 1 hour at room temperature in either 5% bovine serum albumin (BSA) or 4% skim milk prepared in Tris-buffered saline containing 0.1% Tween-20 (TBS-T), followed by incubation with primary antibodies at a dilution of 1:1000 overnight at 4 °C. Primary antibodies included anti-AR (EP670Y; ab52615, Abcam), anti-HaloTag (G9211, Promega), anti-V5 (SV5-Pk1; R960-25, Thermo Fisher Scientific), anti-JunD (EPR17365; ab181615, Abcam), anti-c-Jun (60A8; #9165, Cell Signaling Technology), anti-phospho-c-Jun (Ser63; Y172; ab32385, Abcam), anti-c-Fos (EPR20769, Abcam), anti-NF-κB2 p100/p52 (#4882, Cell Signaling Technology), anti-vinculin (V9131, Sigma-Aldrich), and anti-GAPDH (3E12; bsm-0978M-TR, Bioss). HRP-conjugated anti-β-tubulin (EPR1330; ab185057, Abcam) was also used as a loading control. Following primary antibody incubation, membranes were incubated with the appropriate HRP-conjugated secondary antibody at a 1:5000 dilution: anti-mouse IgG-HRP (62-6520, Invitrogen) or anti-rabbit IgG-HRP (31460, Invitrogen). Protein signals were detected using Pierce™ ECL Western Blotting Substrate (Thermo Fisher Scientific). Immunoblots were quantified by densitometric analysis using ImageJ. Band intensities were normalized to the corresponding loading control and subsequently expressed relative to a control sample included on each blot.

### Chromatin Immunoprecipitation and Sequencing

ChIP was performed on human myoblasts stably expressing AR 24Q-V5 or AR 100Q-V5 under a doxycycline-inducible promoter. Cells were treated with 1 µg/mL doxycycline and 10 nM R1881 or vehicle in FBS-free growth media for 48 hours. For each condition, 5x10 cells were cross-linked with 1% formaldehyde for 10 minutes, then harvested and lysed. Cell lysates underwent sonication (12 cycles of 30 seconds on/off) to achieve DNA fragment lengths of 200-500 bp. Immunoprecipitation was conducted using 14 µg of anti-V5 antibody (SV5-Pk1; R960-25, Thermo Fisher Scientific) or anti-IgG (5415, Cell Signalling Technology) as previously described^107^. DNA samples were purified using the MinElute PCR Purification Kit (Qiagen) to enrich for target fragment sizes and subjected to quantitative real-time PCR. Two biological replicates were included per condition.

Library preparation and next-generation sequencing (NGS) were conducted by Novogene Co., Ltd. (Cambridge, UK). DNA fragments were end-repaired, A-tailed, and ligated with Illumina adapters. The resulting libraries underwent size selection and PCR amplification to generate the final DNA libraries. Following quantification, the libraries were pooled and sequenced using Illumina platforms according to standard protocols.

### ChIP-Seq Analysis

Raw sequencing reads were aligned to the human reference genome (GRCh38) using a BWA and duplicate reads were removed with Picard MarkDuplicates (v3.3.0; https://broadinstitute.github.io/picard). Peaks were called using HOMER (v5.1 http://homer.ucsd.edu/homer/ngs/peaks.html) which identifies significant regions of enrichment. To assess reproducibility between biological replicates, an irreproducible discovery rate analysis was performed using a Python-based implementation (https://github.com/nboley/idr) retaining high-confidence peaks (irreproducible discovery rate <0.05) for downstream analysis. A consensus peak set was generated across samples, and differential binding analysis was conducted using DiffBind (v3.14.0; https://www.bioconductor.org/packages//2.11/bioc/html/DiffBind.html) to identify regions exhibiting significantly altered AR occupancy between experimental conditions (*P* < 0.05). Peak annotation to nearby genes and functional genomic elements was performed with ChIPSeeker (v1.40.0; https://bioconductor.org/packages/release/bioc/vignettes/ChIPseeker/inst/doc/ChIPseeker.html).The MEME suite was used to identify enriched protein motifs (https://meme-suite.org/meme). ChIP-seq signal tracks and genomic regions of interest were visualized using Integrative Genomics Viewer (IGV; https://igv.org/).

Identification of AR-associated super-enhancers and overlap with ALM-associated variants Super-enhancers were identified using the Rank Ordering of Super-Enhancers (ROSE) algorithm (v0.1; https://hpc.nih.gov/apps/ROSE.html). AR ChIP-seq peaks from wild-type cells following R1881 treatment were used as input to ROSE. ROSE was run using default parameters, with using distal and intragenic peaks as input gtf as list of enhancers. The resulting stitched regions were ranked according to AR ChIP-seq signal, and super-enhancers were defined based on the inflection point of the resulting signal-versus-rank curve.

To identify AR-associated super-enhancers overlapping regulatory regions in human skeletal muscle, ROSE-defined super-enhancers were intersected with publicly available histone ChIP-seq data from human psoas muscle tissue generated by the ENCODE Project^108^ (accession ENCSR791ISZ). Only super-enhancer regions exhibiting at least 1,000 bp of overlap with an ENCODE histone ChIP-seq peak were retained for subsequent analyses. To determine whether these super-enhancers contained genetic variants associated with lean muscle mass, the retained regions were compared with the 1,059 conditionally independent single-nucleotide polymorphisms (SNPs) associated with appendicular lean mass (ALM) identified by Pei et al., 2020^49^. As the GWAS SNP coordinates were reported using the hg19/GRCh37 human genome assembly, SNP coordinates were converted to hg38/GRCh38 using the liftOver function from the Bioconductor rtracklayer package and an hg19-to-hg38 chain file (v1.72.0; https://bioconductor.org/packages/release/bioc/html/rtracklayer.html). Only SNPs mapping uniquely to a single location in hg38 were retained; SNPs that failed to map or mapped to multiple genomic locations were excluded. A SNP was considered to overlap a super-enhancer when its hg38 genomic coordinate fell within the start and end coordinates of the super-enhancer.

A regional association plot for the JUND locus was generated from the GWAS summary statistics^49^ using locuszoomr (v0.3.8; https://cran.r-project.org/web/packages/locuszoomr/vignettes/locuszoomr.html). The region spanning ±2 Mb around JUND was plotted, with rs10686842 selected as the index variant. Linkage disequilibrium (LD) was obtained using LDlink, and variants were labelled according to their pairwise LD (R²) with rs10686842. R² values range from 0 to 1, with values approaching 1 indicating strong LD with the index variant and values approaching 0 indicating little or no LD.

### RNA sequencing and differential gene expression analysis

RNA sequencing (RNA-seq) was performed on cells expressing AR 24Q or AR 100Q, cultured in the presence or absence of androgen, with three biological replicates per condition (*n* = 3). Adopting the bioinformatic pipeline (https://nf-co.re/rnaseq/3.14.0/), sequence reads were adapter- and quality-trimmed using Trim Galore! (v0.3.1; www.github.com/FelixKrueger/TrimGalore), with quality control performed using FastQC (v0.11.1; www.bioinformatics.babraham.ac.uk/projects/fastqc/) and MultiQC (v0.7)^109^. Trimmed reads were aligned to the human reference genome (Ensembl GRCh38) using HISAT2 (v2.1.0; https://daehwankimlab.github.io/hisat2), alignment files were processed using SAMtools (v1.3; https://www.htslib.org/doc/1.3.1/samtools.html), and gene-level read counts were generated against the Homo_sapiens.GRCh38.86.gtf annotation using HTSeq (v0.9.1; https://htseq.readthedocs.io/en/release_0.9.1) with htseq-count in union mode.

Differential gene expression analysis was performed in R using DESeq2 (v1.22.2; https://bioconductor.org/packages/release/bioc/html/DESeq2.html). Lowly expressed genes were excluded by requiring ≥10 reads in at least three samples and a maximum count of ≥ 20 reads in at least one sample. Counts were normalized using DESeq2 size-factor normalization, and sample quality and clustering were assessed using relative log expression, principal component analysis of variance-stabilizing transformed counts, and sample-to-sample distances. Differential expression was assessed using Wald tests, with Cook’s distance used for outlier detection and log2 fold changes shrunk using apeglm (v1.34.0; https://bioconductor.org/packages/release/bioc/html/apeglm.html). Pairwise differential expression analyses were performed to assess the transcriptional response to androgen stimulation (AR 24Q -R1881 vs. AR 24Q +R1881) and the effect of AR polyglutamine expansion following androgen stimulation (AR 24Q +R1881 vs. AR 100Q +R1881). *P* values were adjusted for multiple testing using the Benjamini–Hochberg method. A cutoff of *P*adj < 0.01 was used for reported DEGs. Gene set enrichment analysis^48^ was performed using the GSEA preranked module in the Broad Institute GSEA Desktop software. Genes were ranked by the DESeq2 Wald statistic and tested against an AP-1 TF gene set using a weighted enrichment statistic and 1,000 gene-set permutations.

### MRI data acquisition

This prospective cohort study involved UK Biobank participants enrolled in the imaging sub study between January 2016 and February 2020. Individuals aged 40 to 69 years old were invited for MRI examination. All participants had been scanned at one of the UK Biobank imaging centres, on a Siemens Aera 1.5 T scanner (Siemens Healthineers), with both a dual-echo Dixon VIBE protocol for body composition and the LiverMultiScan imaging protocol that are encompassed within the UK Biobank imaging protocol^64,110^. This research was conducted using data obtained using UK Biobank Access Application number 9914. Ethics approval for participation in UKB was obtained from the North West Multi-Centre Research Ethics Committee and all participants provided written informed consent to provide access to their healthcare records. Body composition measurements were derived using fully automated volumetric analyses, as described previously^68^. Volumes for VAT, SAT, abdominal SM volume and MFI were calculated based on the number of voxels. All analyses were performed by trained MR analysists blinded to all clinical data. Liver MRI data analysis was performed automatically using LiverMultiScan® software (Perspectum, Oxford, UK)^111^, and every case was manually reviewed by trained analysts, blinded to all clinical data.

### AR CAG repeat length

The length of the CAG trinucleotide repeat within exon 1 of the *AR* gene was determined from whole-genome sequencing data generated by the UK Biobank. Short tandem repeat (STR) genotypes were obtained from ExpansionHunter^69^ calls provided as part of the UK Biobank DRAGEN whole-genome sequencing pipeline. Variant call format (VCF) files containing the AR CAG repeat locus were extracted and merged across participants using bcftools^112^. ExpansionHunter repeat copy number estimates were extracted for each participant and linked to imaging-derived phenotypes. Participants without valid repeat length estimates or relevant imaging data were excluded from downstream analyses.

### ASO Design and Synthesis

Gapmer antisense oligonucleotides (ASOs) targeting Eda2r were selected using OligoAI, an artificial intelligence model developed to predict ASO *in vitro* knockdown efficacy^87^. The highest-scoring ASOs were transfected into myoblasts at a final concentration of 100 nM using Lipofectamine™ RNAiMAX (Thermo Fisher Scientific). ASO-mediated knock-down of *Eda2r* was subsequently assessed by RT-qPCR.

ASOs were either purchased from Integrated DNA Technologies (IDT; Leuven, Belgium) or synthesized in-house. Solid-phase synthesis of antisense oligonucleotides (ASOs) was performed using a K&A H-16 synthesizer (Sierra BioSystems). DNA and 2′-*O*-MOE phosphoramidites were obtained from ChemGenes, and all cytidine residues in the DNA and 2′-*O*-MOE regions were incorporated as 5-methylcytidine (5-Me-C). All phosphoramidites were dissolved in anhydrous acetonitrile (ACN) to a final concentration of 0.1 M. Detritylation was performed using 3% (v/v) trichloroacetic acid in DCM (Sigma-Aldrich). A 0.3 M solution of Hyacinth 5-benzylmercaptotetrazole (BMT; EMP, supplied by Atom Scientific Ltd) in anhydrous acetonitrile was used as the activator during phosphoramidite coupling. Sulfurization was carried out using 0.1 M DDTT (ChemGenes) in anhydrous pyridine (Sigma-Aldrich), while oxidation was performed using Hyacinth Oxidizer (EMP, supplied by Atom Scientific Ltd), consisting of 0.05 M iodine in a 9:1 (v/v) mixture of pyridine and water. Remaining 5′-hydroxyl groups were capped using Capping A (EMP, supplied by Atom Scientific Ltd), comprising an 8:1:1 (v/v/v) mixture of tetrahydrofuran (THF), lutidine, and acetic anhydride, in combination with Capping B (EMP, supplied by Atom Scientific Ltd), a 16% (w/v) solution of N-methylimidazole in THF. Deprotection and cleavage from the solid support were achieved using 30% aqueous ammonium hydroxide (NH_4_OH; Sigma-Aldrich) at 55°C for 18 h. Afterwards, the solution was allowed to cool to room temperature before the remaining ammonia was removed under reduced pressure using a Speedvac concentrator. Following the deprotection procedure, the oligonucleotides were precipitated in Ethanol puriss (Sigma-Aldrich) and in a 0.3 M Sodium acetate buffer (pH 5.2; ThermoFisher) at -20 °C for 18 h. The precipitated oligonucleotide formed a pellet after the samples were centrifuged at 4550 rpm for 30 min at 4 °C. The resulting pellet was washed twice with 20 mL ice cold 70% ethanol, followed by centrifugation at 4550 rpm for 30 min at 4 °C. Finally, the pellets were dried under reduced pressure, and oligonucleotides were resuspended in RNase-free water (ThermoFisher). If required, additional desalting was achieved using 0.5 mL 3 kDa Amicon Ultra centrifugal filter units (Merck). Liquid chromatography–mass spectrometry (LC-MS) analysis was performed using an ACQUITY Premier UPLC system coupled to a single quadrupole mass detector (SQD2; Waters). Purified oligonucleotides were diluted to a final concentration of 20 µM in water, and 50 µL aliquots were analysed in negative ionisation mode. Separation was achieved using an ACQUITY UPLC BEH C18 column (130 Å, 1.7 µm, 2.1 × 50 mm; Waters). Eluent A consisted of 8.6 mM triethylamine (TEA) and 100 mM 1,1,1,3,3,3-hexafluoroisopropanol (HFIP) in water, while eluent B was methanol. The flow rate was 0.5 mL min⁻¹, and the column temperature was maintained at 60 °C. The mass spectrometer was operated with a capillary voltage of 0.8 kV, a cone voltage of 20 V, and a desolvation temperature of 600 °C. Raw continuum data were deconvoluted to zero-charge mass spectra using MaxEnt1 software (Waters).

### Mice

Animal experiments were performed in the Biomedical Sciences Unit at the University of Oxford, according to procedures authorized by the UK. Home Office (Animal Scientific Procedures Act 1986). Mice were housed in individually ventilated caging systems, with access to food and water ad libitum. AR 100Q transgenic animals^86^ were kindly provided by the Pennuto laboratory. Only male mice were used in the analysis. Animals were aged to 6 weeks prior to initiation of treatment and assigned to treatment groups (*n* = 5 per group). Body weight was recorded prior to treatment initiation and immediately before each subsequent injection. Beginning at 6 weeks of age, animals received intravenous tail-vein injections of either an *Eda2r* targeting antisense oligonucleotide (ASO) or a non-targeting control (NTC) at a dose of 30 mg/kg twice weekly.

### Serum EDA2R quantification

Circulating EDA2R concentrations were quantified in mouse serum using the Mouse EDA2R ELISA Kit (A303585; Antibodies.com) according to the manufacturer’s instructions. Serum samples were diluted 1:4 in sample dilution buffer to ensure measurements fell within the detectable range of the standard curve. All samples and standards were performed in duplicate. Absorbance was measured at 450nm using a microplate spectrophotometer (CLARIOstar, BMG LabTech). EDA2R concentrations were calculated from the standard curve and corrected for the sample dilution factor.

### Immunohistochemistry

Tibialis anterior muscles were fixed in 4% paraformaldehyde for 24 hours. Following fixation, samples were processed using a Tissue-Tek® VIP® 6 automated tissue processor (Sakura Finetek), during which tissues were dehydrated, cleared, and infiltrated with paraffin.

Paraffin-embedded tissues were sectioned at 5 µm using a rotary microtome. Sections were subsequently stained with hematoxylin and eosin (H&E) using a Tissue-Tek® DRS™ 2000 AutoStainer (Sakura Finetek). Whole-slide images were acquired using a ZEISS Axioscan 7 slide scanner with a 20x objective.

### Measurement of nascent protein synthesis

Protein synthesis was assessed using the Click-iT™ Plus OPP Alexa Fluor™ 488 Protein Synthesis Assay Kit (Thermo Fisher Scientific). Myoblasts were seeded and treated with the indicated compounds for 24 hours, followed by incubation with 20 µM O-propargyl-puromycin (OPP) for 30 minutes. Cells were subsequently processed according to the manufacturer’s instructions, and incorporated OPP was detected by Click-iT-mediated conjugation to Alexa Fluor 488. Cells were imaged using an ImageXpress® confocal imaging system (Molecular Devices) with a 20x objective, acquiring five fields per well. Alexa Fluor 488 fluorescence intensity was quantified as a measure of nascent protein synthesis. Cycloheximide (CHX) was used as a negative control.

### Statistical Analysis

A two-tailed Student’s t-test and an ANOVA test with Tukey’s multiple-comparisons correction were used to compare the means between two and three or more groups, respectively. Associations between AR CAG repeat length and imaging-derived phenotypes were evaluated using multivariable linear regression. AR CAG repeat length was modelled as a continuous predictor, such that the regression coefficient represents the expected change in the imaging phenotype per additional CAG repeat. GraphPad Prism (v10; GraphPad, La Jolla, CA) was used for statistical analysis.

## Data Availability

The sequencing data generated in this study have been deposited in the Gene Expression Omnibus (GEO) database. Accession information will be provided upon request and availability.

## Acknowledgements

Tissue processing and histological stains were performed by the histology core facility at the Kennedy Institute of Rheumatology, University of Oxford.

