## Supplementary Figures for "The androgen receptor polyglutamine tract sets the transcriptional balance between muscle growth and atrophy"

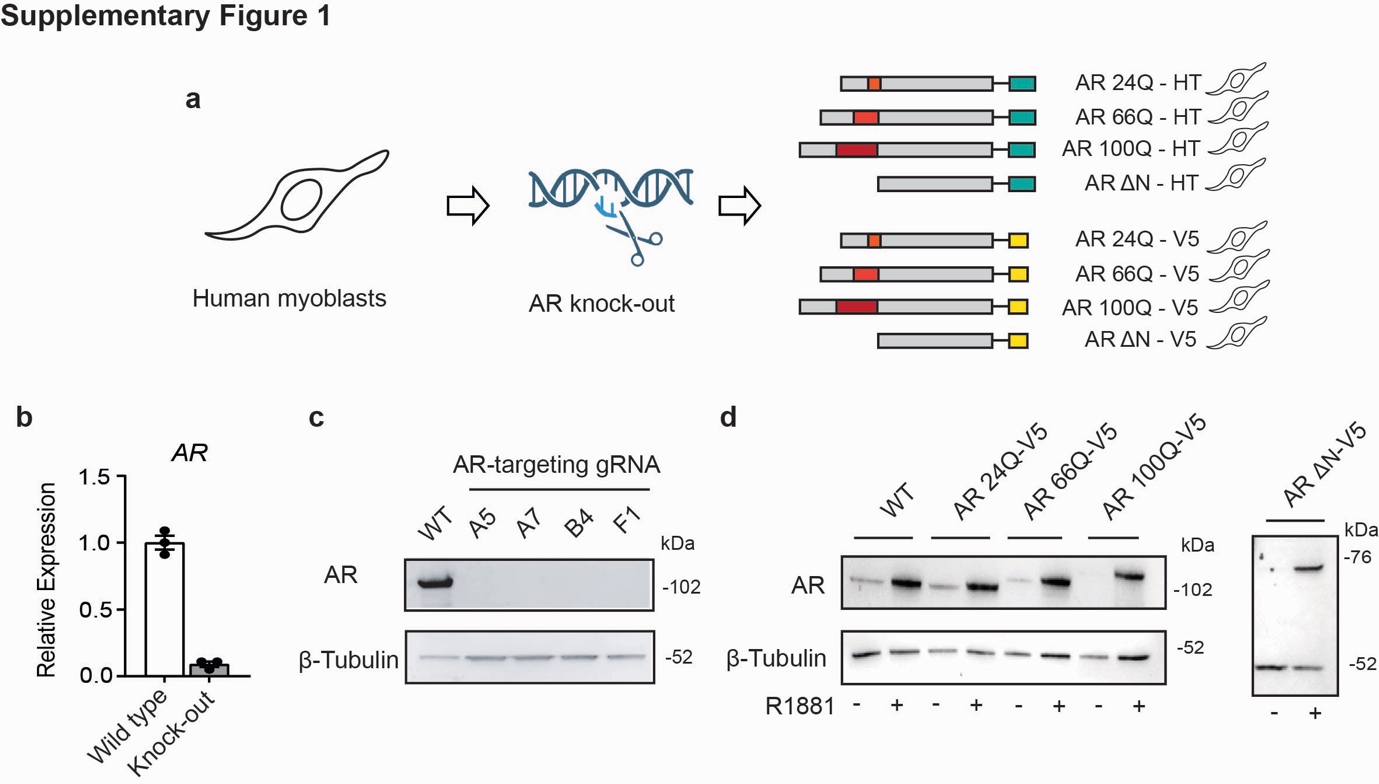

**Supplementary Figure 1. Generation and validation of an isogenic AR allelic series in human myoblasts. a**, Schematic of the experimental system: parental human myoblasts were rendered AR-null by CRISPR–Cas9 and reconstituted with an isogenic allelic series (AR 24Q, AR 66Q, AR 100Q and N-terminally deleted AR ΔN), each tagged at the C terminus with either HaloTag (HT, teal; for super-resolution microscopy and single-particle tracking) or a V5 epitope (yellow; for ChIP). Red block, polyglutamine tract. **b**, Quantitative RT-PCR of *AR* mRNA in parental wild-type versus *AR* knock-out human myoblasts, confirming loss of endogenous AR expression. Data are mean ± s.e.m. (*n* = 3 biological replicates); two-tailed unpaired t-test. **c**, Western blot of AR protein in wild-type (WT) cells and four independent AR-targeting gRNA-edited clones (A5, A7, B4, F1); β-Tubulin, loading control. No residual AR protein is detected in any edited clone. **d**, Western blot confirming re-expression of AR 24Q, AR 66Q, AR 100Q (panel on the right) and AR ΔN (panel on the left), tagged with V5 in the knock-out background compared to the parental cell lines expressing endogenous AR, referred here as wild type (WT). Cells were treated with doxycycline 1 μg/mL to induce transgene expression; β-Tubulin, loading control.

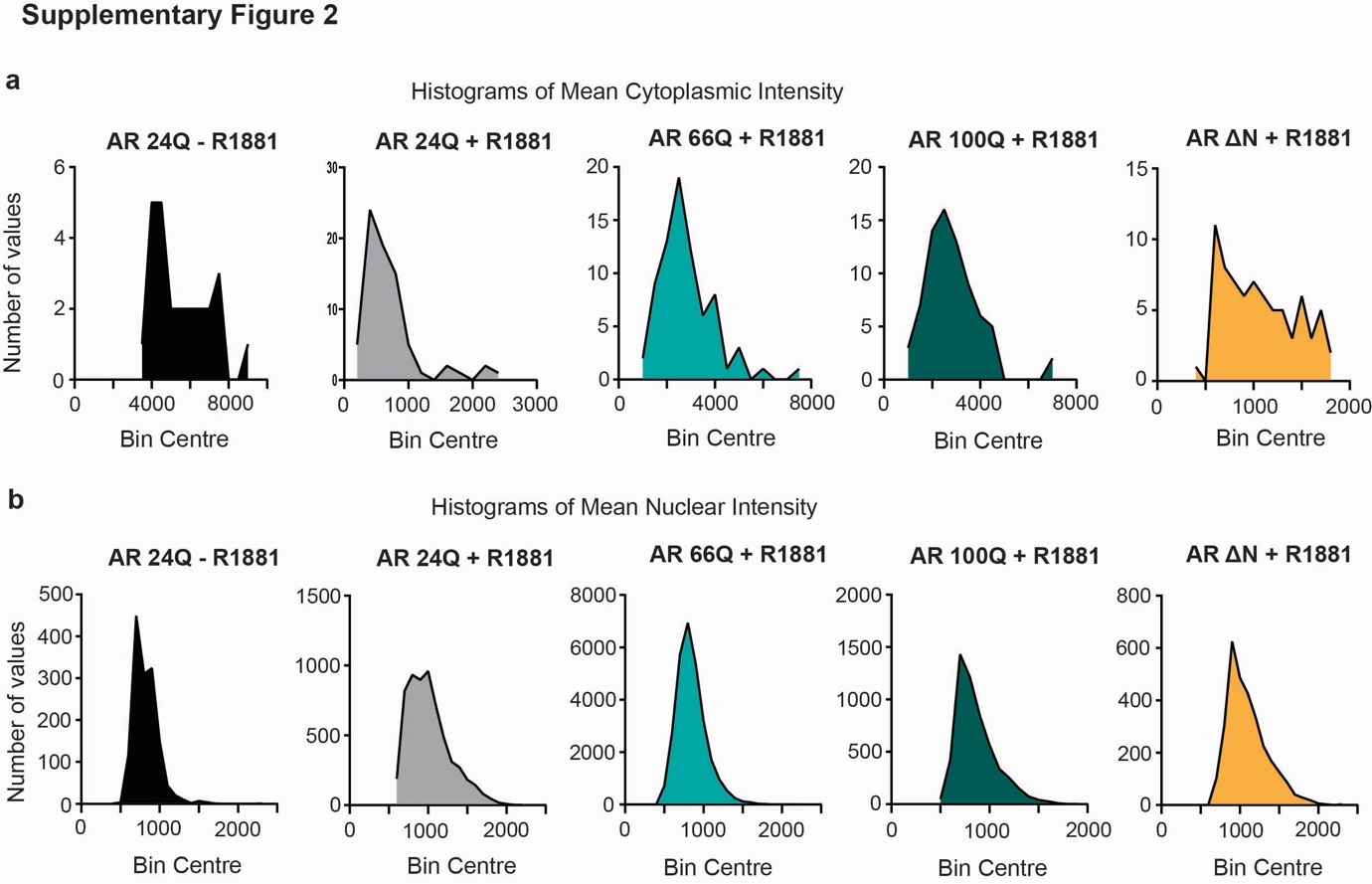

**Supplementary Figure 2. Distribution of cytoplasmic and nuclear AR intensity across the allelic series. a**, Histograms of mean cytoplasmic fluorescence intensity per cell for AR 24Q (-R1881 and +R1881), AR 66Q (+R1881), AR 100Q (+R1881) and AR ΔN (+R1881). **b**, Corresponding histograms of mean nuclear fluorescence intensity for the same conditions. y-axis, number of cells (values); x-axis, binned mean intensity (a.u.).

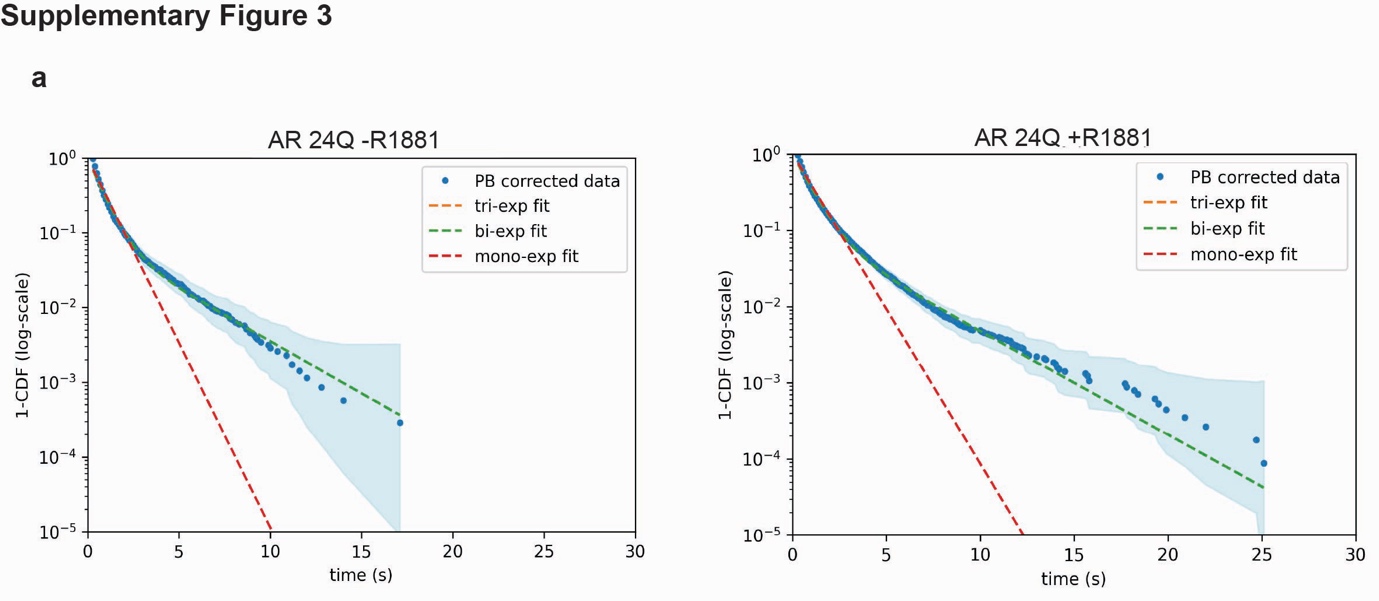

**Supplementary Figure 3. Dwell-time survival analysis of single-molecule AR binding events. a**, Exemplary survival distributions (1 − CDF of dwell times, log scale) of chromatin-bound AR molecules obtained by single-particle tracking for AR 24Q, in the absence (left) and presence (right) of R1881. Blue points, photobleaching (PB)-corrected data; dashed lines, mono-exponential (red), bi-exponential (green) and tri-exponential (orange) fits; shaded band, (95% confidence interval / s.d. across cells). The bi-/tri-exponential fits, but not the mono-exponential, capture the long-lived bound fraction, supporting a multi-state binding model.

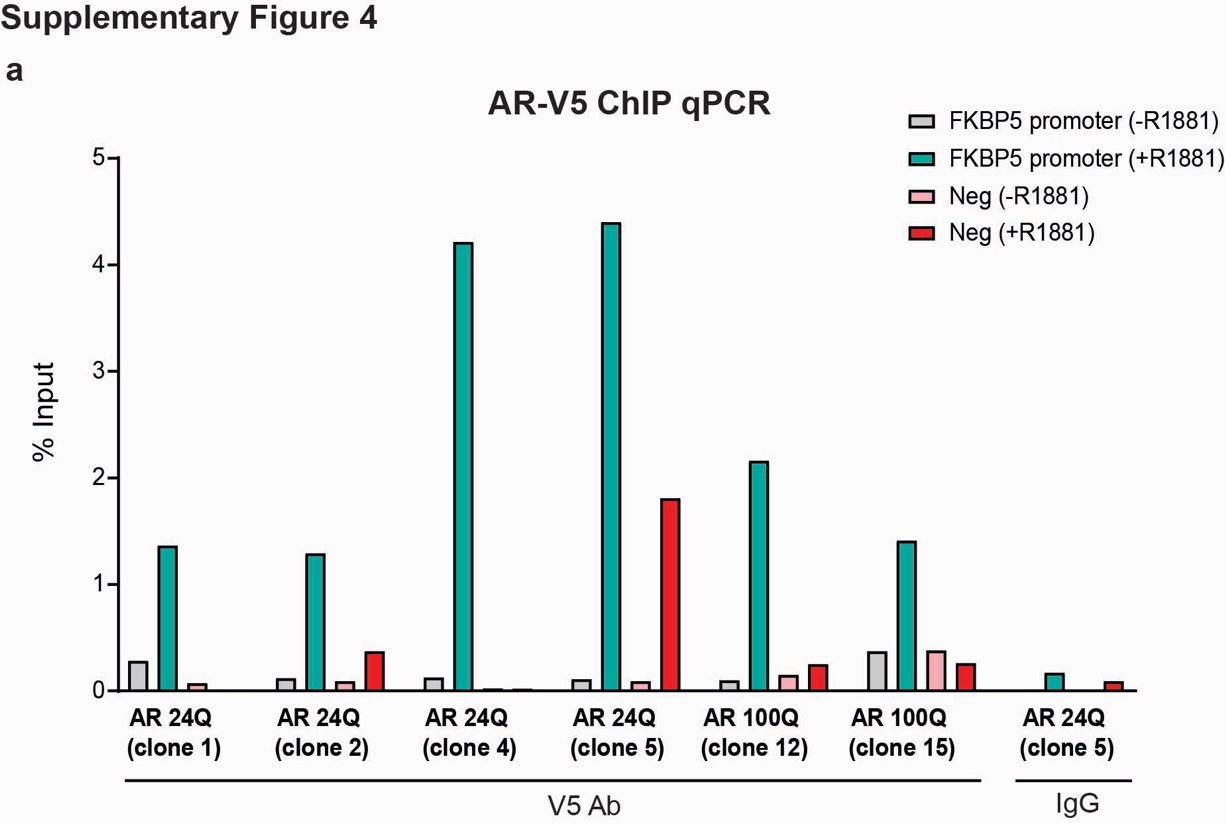

**Supplementary Figure 4. Validation of AR-V5 ChIP by site-specific qPCR. a**, ChIP-qPCR (percentage of input) using an anti-V5 antibody in independent AR 24Q clones (1, 2, 4, 5) and AR 100Q clones (12, 15), assayed at the androgen-responsive *FKBP5* promoter and a negative-control region, each in the absence and presence of ligand. An IgG immunoprecipitation serves as a specificity control. ‘Neg’ indicates a genomic region used as negative control. R1881 induces strong, selective AR enrichment at the *FKBP5* promoter but not at the negative-control locus or with IgG, confirming assay specificity before genome-wide ChIP-seq.

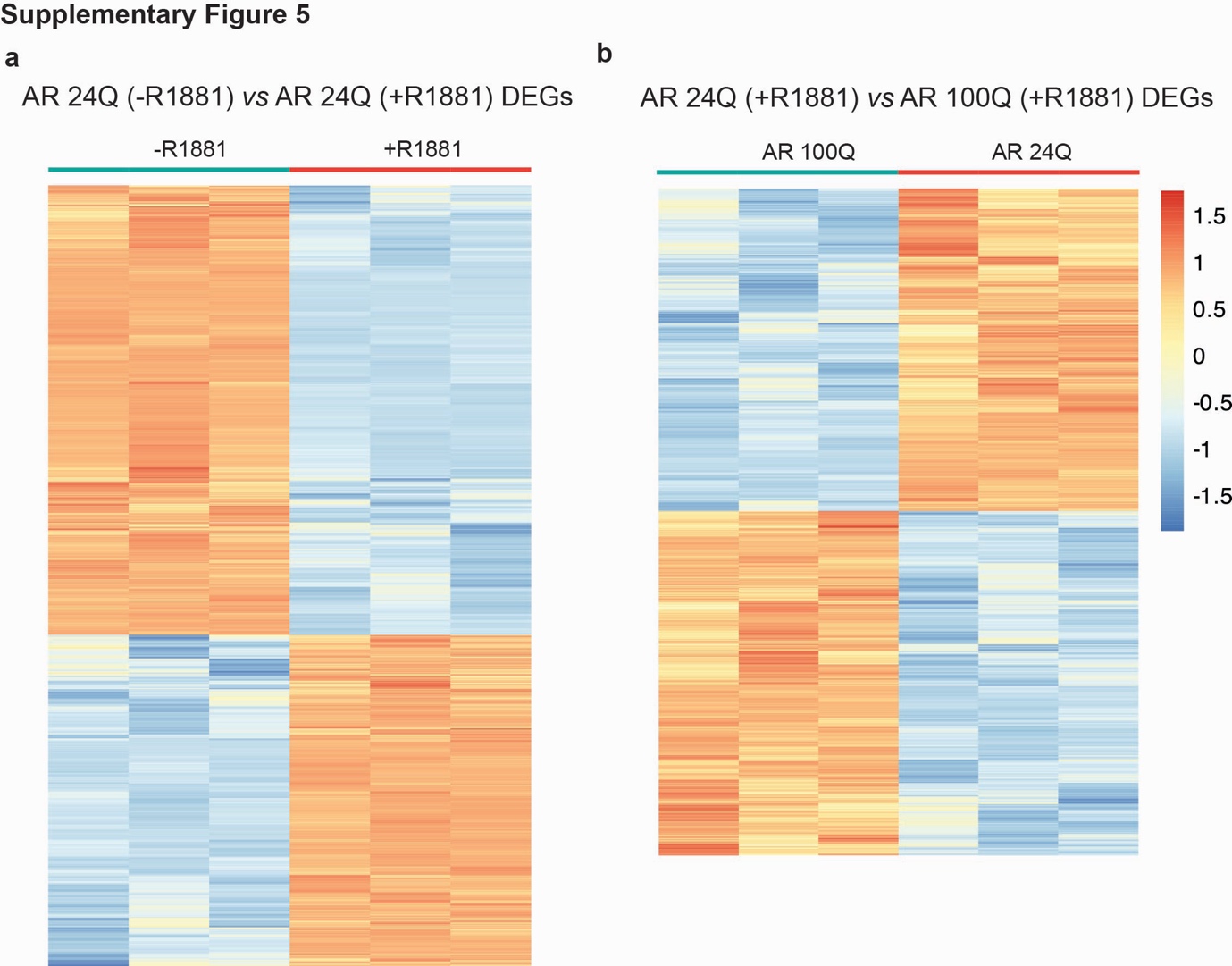

**Supplementary Figure 5. Transcriptional responses to ligand and to polyQ expansion.**

**a**, Heatmap of row-scaled (z-score) expression of differentially expressed genes (DEGs) between AR 24Q -R1881 and AR 24Q +R1881, showing the bidirectional transcriptional response to androgen. **b**, Heatmap of DEGs between DHT-treated AR 24Q and DHT-treated AR 100Q, showing polyQ-dependent transcriptional rewiring. Columns, biological replicates (*n* = 3 per group); colour scale, z-score of normalized expression (-1.5 to +1.5).

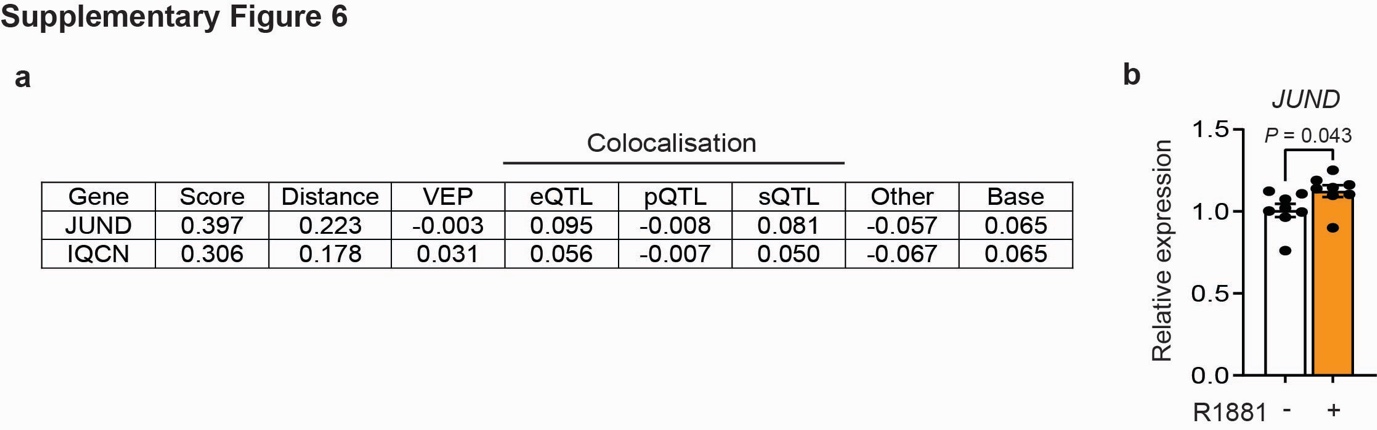

**Supplementary Figure 6. Open Targets locus-to-gene evidence and androgen regulation of JUND. a**, Open Targets Genetics locus-to-gene (L2G) feature table for the two top-ranked candidate genes, *JUND* and *IQCN*, at the rs10686842 locus. Columns show the overall L2G score and its component contributions: distance to canonical TSS, Variant Effect Predictor (VEP) consequence, colocalisation with expression- (eQTL), protein- (pQTL) and splicing- (sQTL) quantitative trait loci, other features, and the model base value. *JUND* is prioritised (L2G = 0.397 versus 0.306 for *IQCN*), driven mainly by TSS proximity together with eQTL and sQTL colocalisation. **b**, Quantitative RT-PCR of JUND in AR 24Q myoblasts without and with R1881, showing modest androgen-induced upregulation. Data are mean ± s.e.m. (*n* = 8); two-tailed t-test, *P* = 0.043.

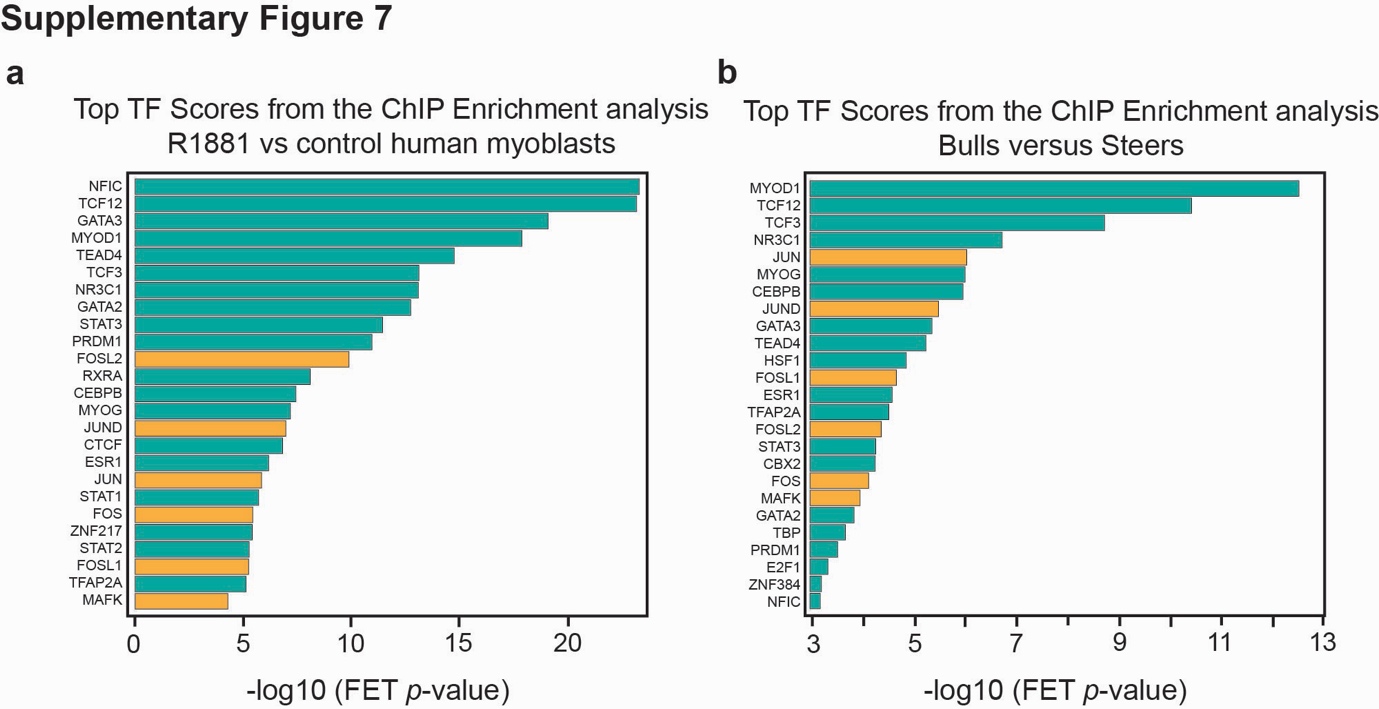

**Supplementary Figure 7. ChIP-X enrichment analysis (ChEA3) of DHT- and androgen-responsive transcriptional programmes. a**, Top transcription factors ranked by ChIP enrichment (−log₁₀ Fisher's exact test *P*-value) for genes differentially expressed in DHT-treated versus control human myoblasts. **b**, Equivalent ChIP-enrichment ranking for genes differentially expressed between bulls and steers (*in vivo* androgen-dependent skeletal-muscle transcriptome; PMID 39633259). AP-1 complex members (FOS, FOSL1, FOSL2, JUN, JUND, MAFK) are highlighted in orange; all other factors in teal. AP-1 components are collectively enriched in both the cell-based and in vivo androgen-responsive programmes.

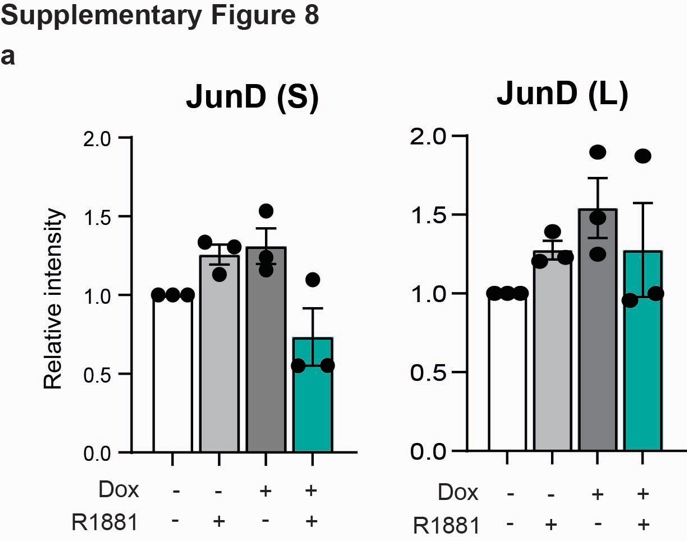

**Supplementary Figure 8. Androgen and doxycycline dependence of JunD isoform induction. a**, Western-blot quantification of the short (JunD (S)) and long (JunD (L)) JunD isoforms across four conditions defined by doxycycline (Dox, AR induction) and DHT treatment. Relative intensity normalized to the -Dox/-R1881 control. Data are mean ± s.e.m. (*n* = 3 independent experiments).

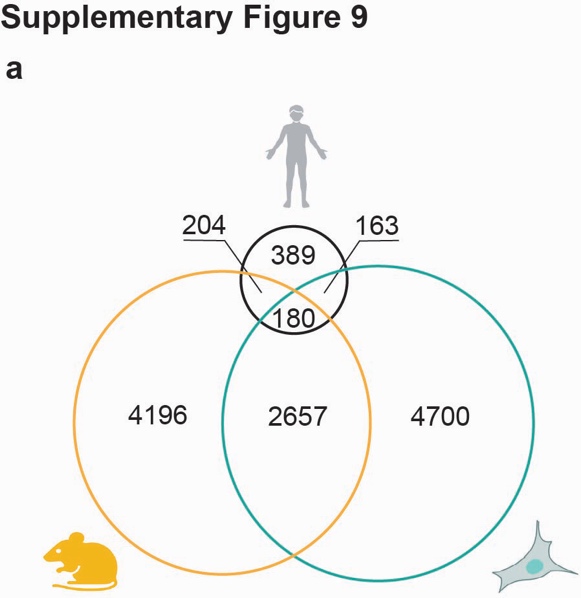

**Supplementary Figure 9. Overlap of androgen/polyQ-responsive transcriptomes across model systems. a**, Three-way comparison of differentially expressed genes between SBMA patient muscle (top, human silhouette), the SBMA mouse model (orange) and AR 100Q human myoblasts (green). Numbers indicate genes unique to or shared between compartments; of the patient-associated DEGs, 389 overlap the patient set, with 204 shared with mouse, 163 with myoblasts and 180 common to all three. The shared core defines a conserved androgen/polyQ-responsive gene set.

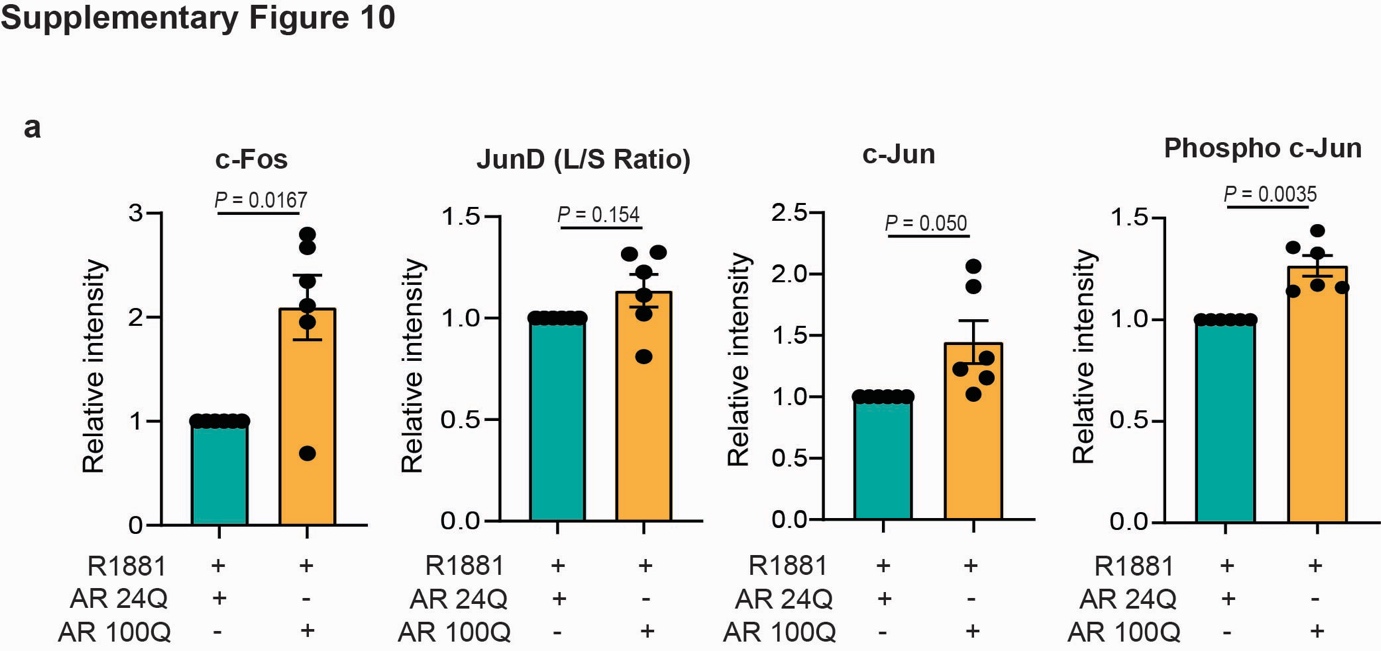

**Supplementary Figure 10. polyQ-dependent activation of AP-1 components under androgen. a**, Western-blot quantification of c-Fos, the JunD long/short (L/S) isoform ratio, c-Jun and phospho-c-Jun in R1881-treated AR 24Q versus DHT-treated AR 100Q myoblasts. Relative intensity normalized to AR 24Q. c-Fos, c-Jun and phospho-c-Jun are significantly higher in AR 100Q; the JunD L/S ratio trends upward. Data are mean ± s.e.m. (*n* = 6); two-tailed t-test, *P*-values as indicated.

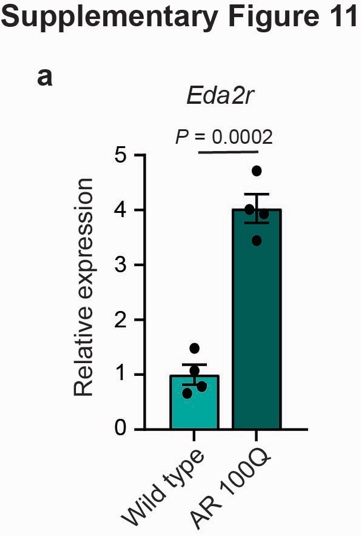

**Supplementary Figure 11. Eda2r is induced in polyQ-expanded muscle. a**, qRT-PCR of *Eda2r* in wild-type versus AR 100Q mice, showing ~4-fold induction in the polyQ-expanded background. Data are mean ± s.e.m. (*n* = 4); two-tailed t-test, *P* = 0.0002.

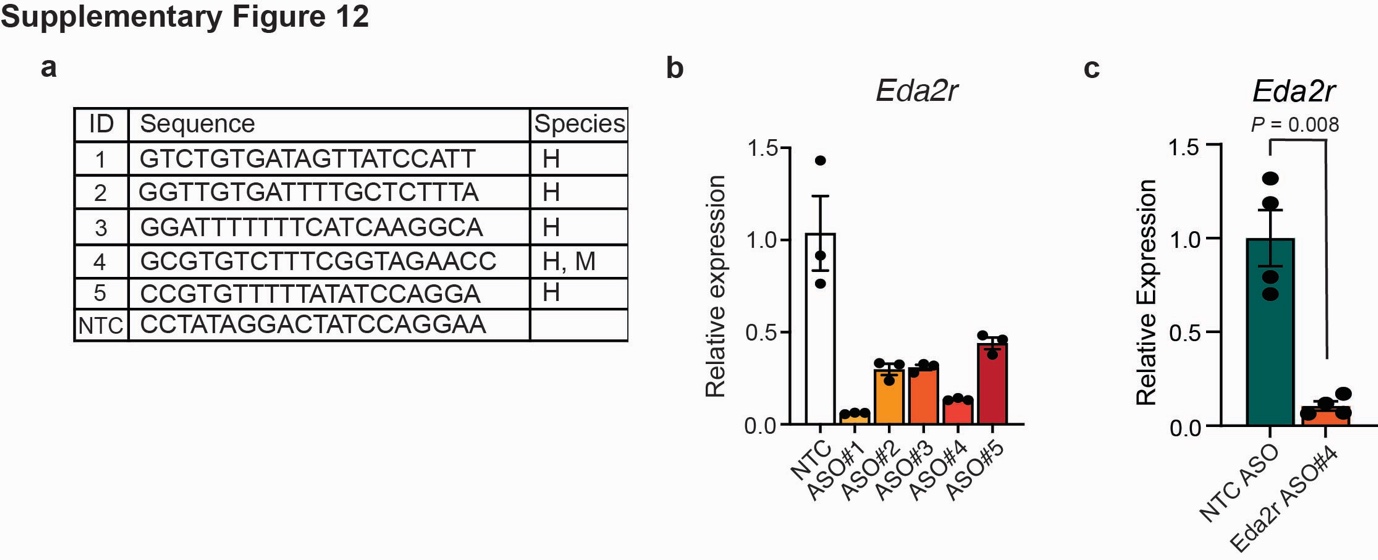

**Supplementary Figure 12. Design and validation of Eda2r-targeting antisense oligonucleotides. a**, Sequences and species specificity (H, human; M, mouse) of the five candidate *Eda2r*-targeting and non targeting control (NTC) gapmer ASOs (phosphorothioate backbone and 2′-MOE wings). **b**, *Eda2r* knock-down efficiency (qRT-PCR) for the five ASOs relative to non-targeting control (NTC) in hMyo, identifying ASO#1 and ASO#4 as most potent. **c**, Validation of the selected human/mouse cross-reactive ASO#4 versus NTC ASO, confirming robust *Eda2r* silencing. Data are mean ± s.e.m. (*n* = 3-4 biological replicates); two-tailed t-test, *P* = 0.008 in c.
